# Identification of APOE4 modulators, targeted therapeutic candidates in Coronary Artery Disease, using Molecular Docking studies

**DOI:** 10.1101/2021.02.02.429307

**Authors:** Lima Hazarika, Supriyo Sen, Akshaykumar Zawar, Jitesh Doshi

## Abstract

A significant genetic suspect for coronary artery disease is the pathological adaptation of apolipoprotein E4 (APOE4) through intramolecular interaction. With the prevailing evidences on APOE4 genotype and its prevalence in coronary artery disease, the present study has investigated the protein–ligand binding affinity and unveil the receptor binding abilities of different classes of ligands for APOE4 through molecular docking studies. Structural basis of APOE4 involvement in CAD suggests that the intramolecular domain interactions to be a suitable target for therapeutic intervention. Various classes of ligands including known drugs used in the treatment of CAD, fragment-based stabilizers and their similar structures and molecules with known bioactivity against APOE4 were screened for their binding affinity and further investigated for their interactions with APOE4. Computational studies show the benzyl amide derived structures to be useful candidates in modulation of APOE4. The dynamics of the binding analysis can be further achieved with an in-depth understanding of drug-receptor interactions performing molecular dynamic simulation studies.

## Introduction

Coronary artery disease (CAD) also known as coronary heart disease (CHD) characterized by cholesterol build up on the inner walls of coronary arteries leading to a process called atherosclerosis. (CDC; NHLBI, NIH). While genetics play an important role in the pathogenesis, it is believed that cholesterol metabolism plays a central role to elevate low density lipoprotein-cholesterol (LDL-C) levels in plasma and eventually leads to the development of CAD (Moriarty, 2009). Alterations in serum lipoproteins levels or accumulation of elevated levels of LDL cholesterol affects the homeostatic control of cholesterol metabolism resulting in atherosclerotic vascular events such as myocardial infarction, stroke, or peripheral vascular occlusion, a strong predisposition to early CHD (Freeman, 2006). Recent study on clinical and coronary angiographic profiles of symptomatic coronary artery disease (CAD) patients less than 30 years of age in Kerala, India reported low levels of high-density lipoprotein (HDL) and high levels of LDL with long-term mortality rates (Gopalakrishnan et al., 2020). Along with abnormal lipid levels, it has been found that APOE4 (E4/E4) genotype has a relationship with myocardial infarction in Indian patients from South Africa and strongly associated with CAD when investigated among coronary angiographed Punjabi population (north west India) (Ranjith et al., 2004; P. Singh et al., 2008). Similar study also reported the association of APOE4 variant with increased LDL levels and total cholesterol in Kashmiri population (Afroze et al., 2016) indicating APOE can be a potential susceptibility locus for CAD.

Apolipoprotein (APOE) encoded by APOE gene, is a plasma glycoprotein of 34.15 kDa with 299-amino acids (Frieden & Garai, 2012) where the receptor-binding region lies in the N-terminal domain (1-164 residues) targeted for fragment-based drug discovery; the C-terminal domain that contains the lipid-binding function runs from 165-299 residues (Petros et al., 2019; Weisgraber, 1994). APOE is associated with HDL, very low density lipoprotein (VLDL), and majorly with chylomicrons (Singh et al., 2006; Frieden et al., 2017) that regulate lipoprotein metabolism and control the transport and redistribution of lipids among tissues and cells through receptor-mediated pathways (Weisgraber, 1994; Freeman, 2006). It’s cholesterol-raising effects in atherosclerosis and premature cardiovascular diseases (CVD) (Lamia et al., 2011) has made a potential genetic marker with profound influence on the risk of developing neurological and cardiovascular disease through cholesterol metabolism (Mahley, 2016).

The human polymorphic APOE gene has single amino acid substitution at 112 and 158 position resulting in three main alleles: epsilon 2 (ε2), epsilon 3 (ε3) and epsilon 4 (ε4), coding for three isoforms: APOE2 (Cys112/Cys158), the most prevalent; APOE3 (Cys112/Arg158) and APOE4 (Arg112/Arg158) (Table 1) with six possible genotypes: E2/E2, E2/E3, E3/E3, E3/E4, E4/E4 and E2/E4 (Weisgraber, 1994; Boulenouar et al., 2013). Differential effect of APOE genotypes have been studied and found that the gene affects the lipoprotein clearance mechanisms and consequently the lipid profile gets disturbed leading to damage to the cardiovascular system. The APOE4 allele has become an independent risk factor that has influence on lipid profiles and is associated with the development of both type 2 diabetes mellitus (T2DM) and CVD (Eichner et al., 2002; El-Lebedy et al., 2016; Liu et al., 2019). The sequence dissimilarity among APOE2, APOE3 and APOE4 cause intramolecular domain interaction due to difference in their isoelectric points; the three isoforms differ sequentially by one charge unit and hence in the binding affinities to LDL receptors and lipoproteins particles (Weisgraber, 1994; Mahley & Rall, 2000). The polymorphism imparts distinct functional effects in lipoprotein metabolism through the hepatic binding, uptake, and catabolism of lipid particles to these events: a) increase intestinal cholesterol absorption, b) affect LDL synthesis in the liver, and c) raised levels of total cholesterol (TC) and low density lipoprotein cholesterol (LDL-C) eventually leading to higher risk for CAD unlike the other isoforms. (Lamia et al., 2011; Karahan et al., 2015; Hone et al., 2019).

**Table: 1.** Summary of the differences among APOE isoforms.

| Differences | APOE4 | APOE3 | APOE2 | Reference |
| --- | --- | --- | --- | --- |
| Sequence differences w.r.t Residues at position | 112: Arginine | 112: Cysteine | 112: Cysteine | (Rall et al., 1982; Rezeli et al., 2015) |
|  | 158: Arginine | 158: Arginine | 158: Cysteine |  |
| Domain Interaction | Ionic Interaction between Arg-61 and Glu-255 | Absent | Absent | (Mahley, 2016) |
| Binding Affinity | APOE4 binds to large LDL and VLDL | APOE3 and APOE2 bind to small HDL |  | (Frieden & Garai, 2012; H. Li et al., 2013; Phillips, 2014) |
| Prevalence (Allelic Frequency) | 14% | 78% | 7% | (Phillips, 2014) |
| Clinical Association | CAD, CVD, AD | Normal | Type-III Hyperlipoproteinemia | (Moriarty, 2009) |

*CAD: Coronary Artery Disease; CVD: Cardiovascular Disease; AD: Alzheimer Disease; E4: Apolipoprotein4; E3: Apolipoprotein E3; E2: Apolipoprotein E2; LDL: Low density lipoprotein; VLDL: Very Low-Density Lipoprotein; HDL: High Density Lipoprotein. Arg: Arginine amino acid; Cys: Cysteine amino acid.*

The distinctive roles of three isoforms in disease pathogenesis was justified by the 112 arginine residue in N-terminal domain that influences the lipid-binding properties of the C-terminal domain, indicating an interaction between the domains and thus affect the receptor-binding activity (Weisgraber, 1994). Substitution at 112 in APOE4 causes the side chain of Arginine at 61 to extend away from the N-terminal domain resulting it to interact ionically with glutamic acid at 255 in C-terminal domain, unlike that in APOE3 and APOE2 which do not exhibit such domain interaction (Mahley, 2016). Studies also claimed that this substitution in APOE4 also prevent disulphide bond formation with APOA-II, an apolipoprotein found on HDL-C particles and thus facilitate the increased binding to VLDL (Moriarty, 2009) during pathogenesis. It has been now considered that high-throughput screening for small molecules as therapeutics can alter the ligand binding region, blocking the domain interactions in APOE4, thereby altering the functional characteristics in such that they would mirror to those of apoE3 (Frieden & Garai, 2012; Mahley, 2016). Therefore, interruption of N-terminal domain interaction (ionic interaction between arginine −61 and glutamic-255) to change the binding preference of APOE4 from LDL, VLDL to HDL (Mahley, 2016); or by affecting lipidation can help APOE4 structural modification to portray APOE3 like molecule (Safieh et al., 2019) can be a suitable approach for CAD therapeutics.

The underlying precise mechanism by which APOE4 contributes to CAD development is not proven till date, however, cholesterol efflux, an important part of reverse cholesterol transport (RCT) pathway, can be considered to play a central position (Ohashi et al., 2005). Mechanisms on binding affinity of APOE to lipid have been kept forward since long but a possible model explaining the isoform difference has not been proposed yet (Frieden et al., 2017) in relation to CAD. The pathological effects of APOE4 in relation to lipid metabolism could be referred to the hypolipidation state opposing the body’s physiological consequences and thus, becomes less effective in cholesterol efflux induction compared to APOE3 (Safieh et al., 2019). However, with prevailing evidences from studies, not a single and important pathway has been identified, so far, for the driving pathological effects of APOE4 in CAD. Studies discussed impairment of cholesterol efflux with APOE4 accumulation in the endosomal compartments of the cells which leads to increase intracellular cholesterol production, owing to its preferentially binding and partition into chylomicron and VLDL. This assist for chylomicron and VLDL catabolism resulting in elevated LDL-cholesterol and therefore adversely accelerate atherosclerosis (Chou et al., 2006) in CAD (Figure1). Chou et al. suggested that APOE4 has more flexibility in retaining a bond with the LDL-R owing to its lack of disulphide bonds (Chou et al., 2006) and the higher affinity for lipid particles forms a complex that inhibit the release of cholesterol, downregulates the hepatic receptor resulting in an elevation of plasma cholesterol (Moriarty, 2009). In diseased conditions when triacylglycerol-rich lipoprotein (TGRL), chylomicron remnants (CR), VLDL, and intermediate density lipoprotein (IDL) increases, it inhibit the cholesterol efflux from macrophages to APOA-I, that blocks the expression of APOE, along with ATP-binding membrane cassette transporter A1 (ABCA1) and, scavenger receptor B1 (SR-B1) proteins. Thus the free cholesterol within the cells and cholesteryl esters stored gets affected, lending as risk factors for coronary artery disease (Ohashi et al., 2005).

**Figure 1:**
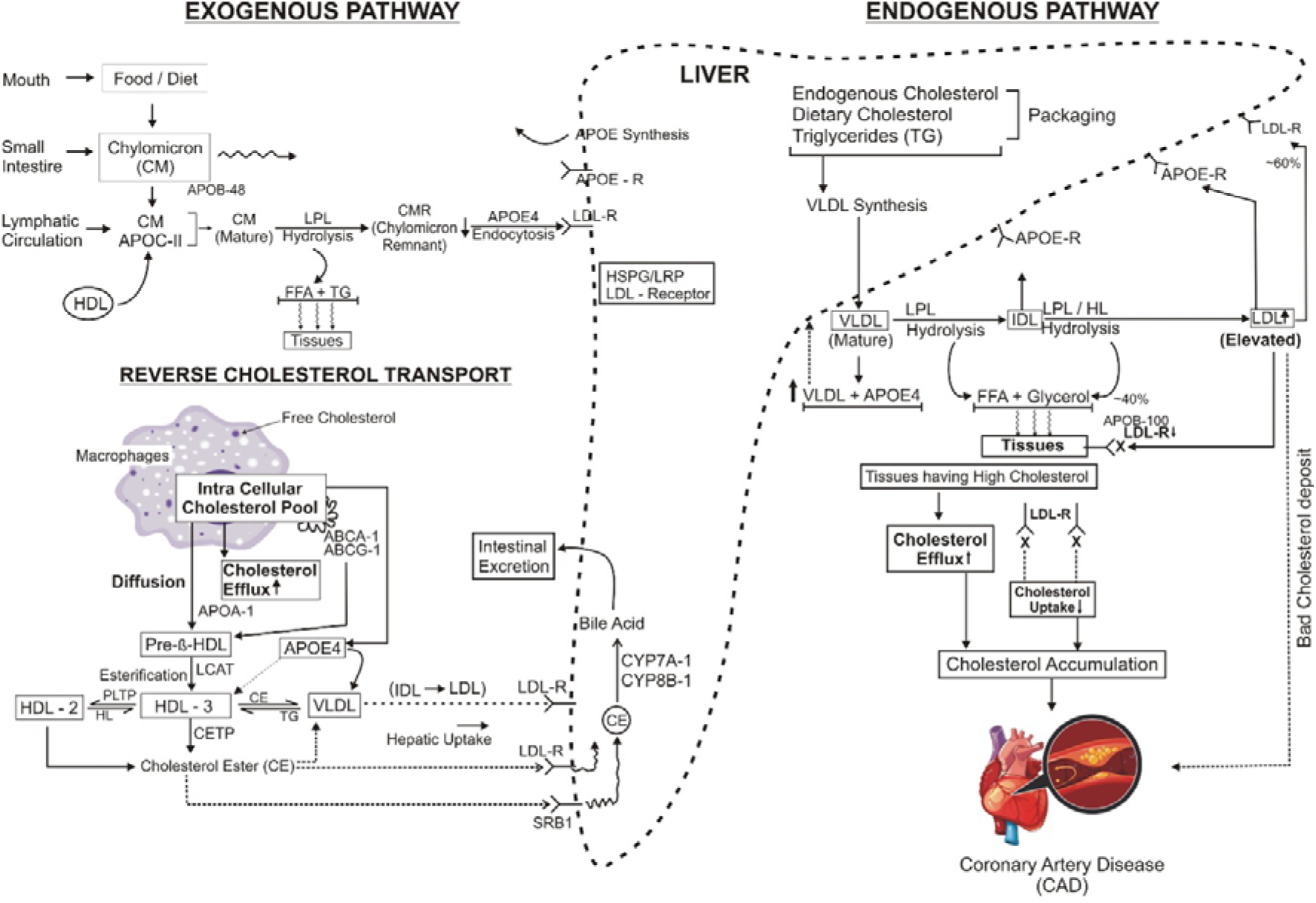
Chylomicron, VLDL and LDL Metabolism in APOE4 isoform leading to impaired cholesterol efflux. The sketch of APOE4 mechanism in the lipoprotein metabolism towards CAD pathogenesis, through three pathways. **The exogenous pathway:** Chylomicrons (CM) are formed in the small intestine after enzymatic digestion of dietary food (triacylglycerol, cholesterol esters, phospholipids, vitamins-D, E, K, A). Entering the lymphatic circulation, they acquire APOE C-II from the circulating HDL to become the matured CM, which undergo hydrolysis by lipoprotein lipase (LPL) and form CM remnants (CMR). The free fatty acids (FFA) and triglycerides (TG) released from the hydrolysis is taken up the peripheral tissues, such as skeletal muscle and adipose tissues. CMR binds to cell surface receptors such as, low density lipoprotein (LDL) receptor (LDLR) or LDLR-related protein (LRP) and heparan sulfate proteoglycan (HSPG) pathways and undergo hepatic clearance. The endocytosis of CMR to Liver is facilitated by APOE receptors. **The endogenous pathway:** Liver synthesize and secretes very low-density lipoproteins (VLDL) which are hydrolyzed by LPL and hepatic lipase (HL) to release of FFA and glycerol, taken up by the tissues. VLDL acquire APOE and APO C-I, II, III from hepatocytes or circulating HDL in blood circulation. Hydrolysis of VLDL results in the formation of intermediate density lipoprotein (IDL) and low-density lipoprotein (LDL) which contain APOB-100, for cellular functions. LDL becomes rich in CE; Under normal conditions, cholesterol regulation of the LDL-R prevents foam cell formation via LDL-R as 60% of matured LDL is taken by LDL-R in liver and 40% of mature LDL is taken by LDL-R to extrahepatic tissues leading to cholesterol accumulation in cells. In pathological conditions, intracellular cholesterol exceeds to release outside the cells called as cholesterol efflux, where, LDL-R expression decreases on the membrane as a result it could not accept or take into anymore LDL. Circulating raised LDL level leads to accumulation of cholesterol in vessels leading to atherosclerosis and CAD. **The reverse cholesterol transport (RCT)**: the pathway through which accumulated cholesterol from peripheral tissues is transported to the liver through high density lipoproteins (HDL) containing APOE. N-HDL triggers cholesterol efflux in macrophages and fibroblasts that absorbs the cholesterol and esterified by LCAT. N-HDL becomes larger resulting in HDL3 (cholesteryl ester rich) and HDL2 (phospholipid rich). PLTP can fuse HDL3 to form HDL2 and HL can process HDL2 and convert to HDL3. CETP facilitates delivery of cholesteryl esters to liver via LDL-R which converts to Bile salts and then eliminate through the GI Tract.

APOE: Apolipoprotein-E; CM: chylomicron; CMR: chylomicron remnant; CE: cholesteryl ester; FFA: free fatty acids; TG: triglycerides; HDL: high density lipoprotein; N-HDL: nascent high density lipoprotein; HL: hepatic lipase; HSPG: heparan sulfate proteoglycan; VLDL: very low density lipoprotein; IDL: intermediate density lipoprotein, LDL: low density lipoprotein, LDL-R: low density lipoprotein receptor; LPL: lipoprotein lipase; LRP: low density lipoprotein receptor-related protein; RCT: reverse cholesterol transport; ER: endoplasmic reticulum; PLTP: phospholipid transfer protein; CETP: cholesterol ester transfer protein; LCAT: Lecithin cholesterol acyltransferase; SRB-1: scavenger receptor, class B type 1.

The distinctive functional aspect of APOE4 owing to its structural difference from the other two isoforms, has created an absolute necessity to understand the underlying interaction of ligand-protein binding and excavate the molecular basis of the disease. Since the APOE4 genotype has been identified as a strong risk factor for CAD, the E4 isoform can be considered as a good target for CAD drug discovery. Till now there is no drug available to inhibit this protein or target the intra domain ionic interaction of APOE4. With very limited reference to the commercially accessible inhibitors for CAD, the present study aimed at identifying the potential therapeutic modulators by structure-based drug discovery (SBDD) method utilizing the 3D structural information of the biological target (Lionta et al., 2014) and discover potential lead drugs for CAD therapy. Computer aided drug design widely used efficient approach for the rapid identification, analysis, and characterization of drug-like candidates in target therapy (Chen, 2013).

## Materials and Methods

### Sequence and Structure Comparisons

The protein sequence of human Apolipoprotein E was retrieved in FASTA format from UniProt (https://www.uniprot.org/) database (UniProtKB -P02649) which is a 317 amino acid long precursor protein. The APOE protein sequence retrieved was analysed for the variants and subjected for sequence comparisons, using Basic Local Alignment Search Tool (BLAST) server of National Centre for Biotechnology Information (NCBI) (https://blast.ncbi.nlm.nih.gov/Blast.cgi) against Protein Data Bank (PDB) database (https://www.rcsb.org/). The protein-protein BLAST analysis used the expected threshold value of 0.05 and the word size 6 with Blosum-62 matrix.

The accession IDs returned from the BLAST analysis were 1LE4, 1B68, 6NCO, 1GS9, 2L7B, 1NFN, 1BZ4, 6CFE, 6NCN. The PDB Ids 2L7B and 1B68 without mutation, showing high percentage of sequence identity, query coverage and the alignment score were selected for further comparison and analysis. The respective sequence and structures of the selected templates were searched in RCSB Protein Data Bank.

Full length sequence of APOE4 was derived from the APOE3 sequence by substituting the amino acids at position 130 (112 in mature protein) and position 176 (158 in actual protein). In literature, APOE protein sequence has been reported with two number schemes: a)1 – 317, which also contains 18 amino acid signal peptides, in which, APOE4 is represented as ARG (130)/ARG (176) and b) 1 – 299, which does not include signal peptide, making APOE4 as ARG (112)/ARG (158). The differences among the APOE isoforms are summarised in the Table 1.

### Homology Modelling

The sequence for APOE4 was subjected for BLAST analysis followed by multiple sequence alignment with 2L7B and 1B68 and eventually to protein model building using Modeller 9.24 (https://salilab.org/modeller/). Homology modelling was performed using UCSF Chimera interface (https://www.cgl.ucsf.edu/chimera/) that enables an interactive visualization and analysis of molecular structures and related data. The final model (Figure 2) was selected based on the lower discrete optimized protein energy (DOPE) score which was −0.24

**Figure2 a:**
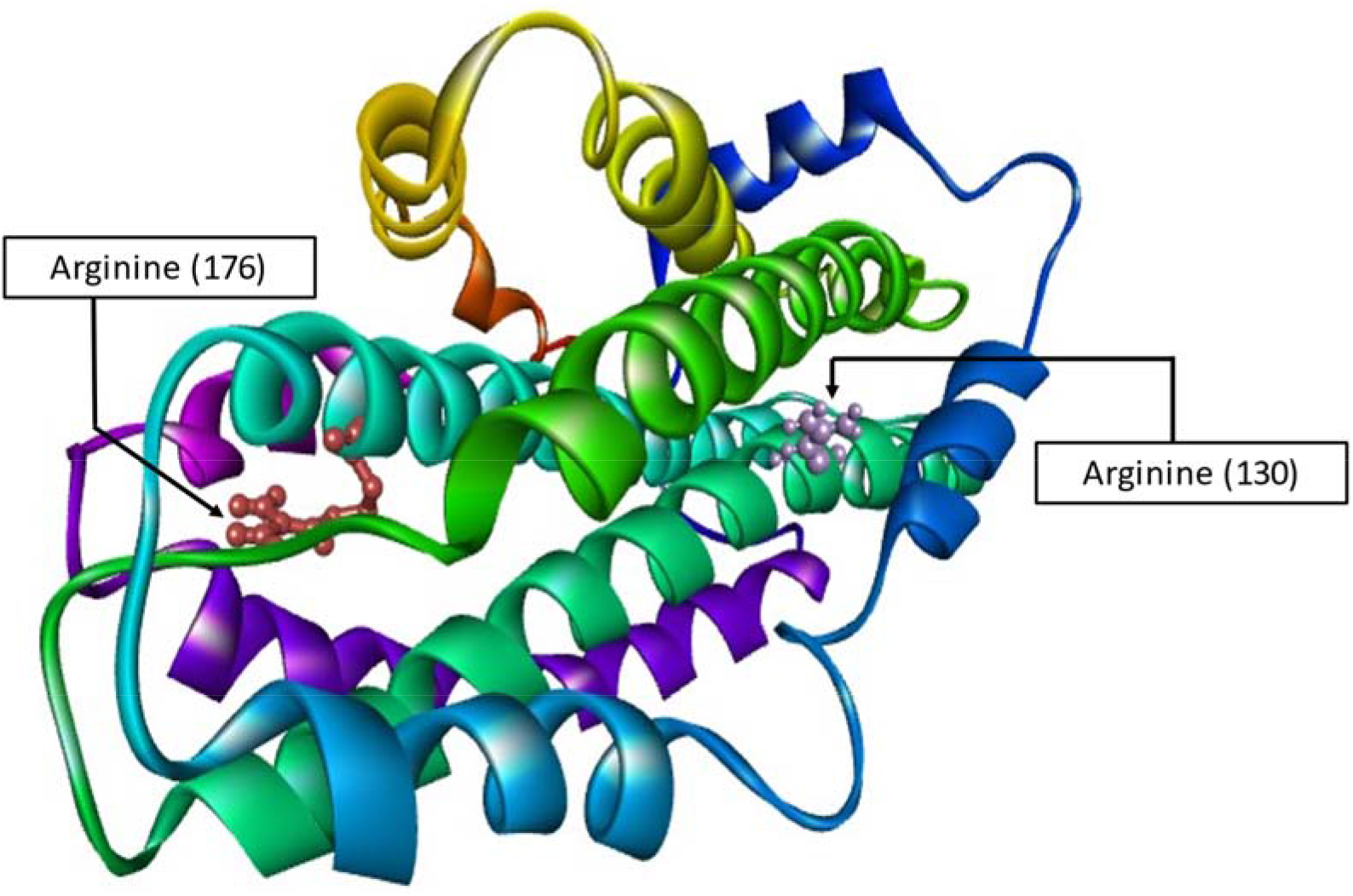
APOE4 Model with zDOPE score as −0.24. Full length sequence of generated APOE4 with modification of Arginine residue at 130 and 176 position, as the sequence retrieved from Uniport consisted first 18 residues for signal peptides. The Arginine residue in actual APOE4 protein is in 112 and 158 respectively.

**Figure2 b:**
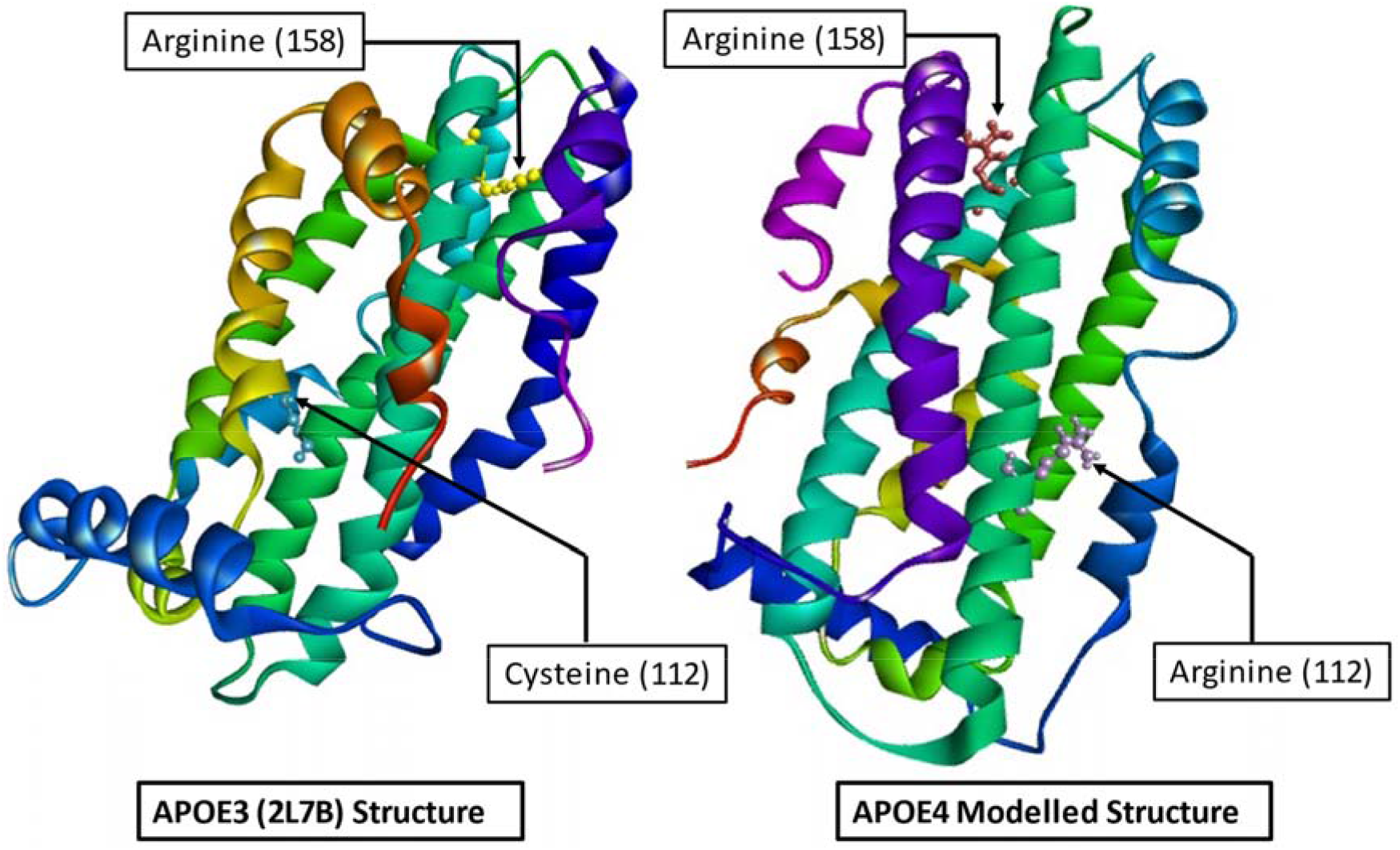
A representation of APOE4 and APOE3 structures.

### Structural assessment

The model generated in UCSF Chimera 1.14 was validated both on geometric and energetic scale using PROCHECK from PDBsum (https://www.ebi.ac.uk/thornton-srv/software/PROCHECK/) and Discovery Studio. This suite of program checks the stereo-chemical properties of the protein, analysed by the Ramachandran Plot, peptide bond planarity, non-bonded interactions, main chain hydrogen bond energy, C-α chirality and overall G factor. Model structure of APOE4 was further minimized using YASARA minimization server (http://www.yasara.org/) in presence of water as solvent to improve the side chain rotations (Krieger et al., 2009). The Ramachandran plot for the APOE4 model before and after energy minimization is shown in Figure 3.

**Figure 3.a:**
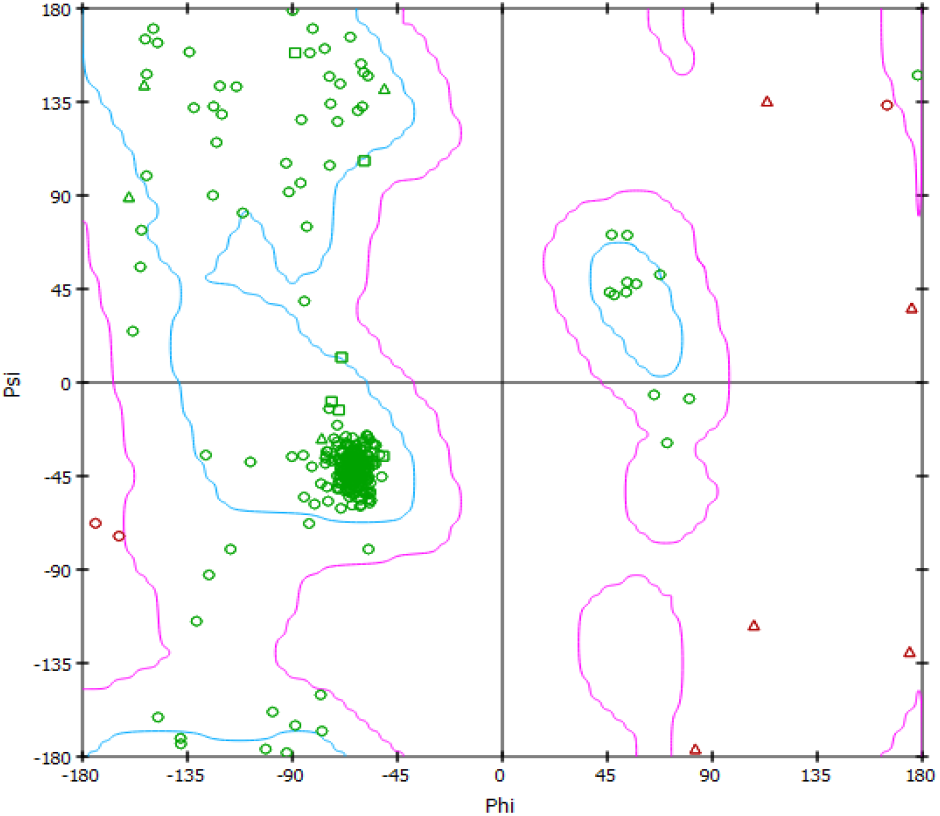
The Ramachandran plot for the APOE4 generated model.

**Figure 3.b:**
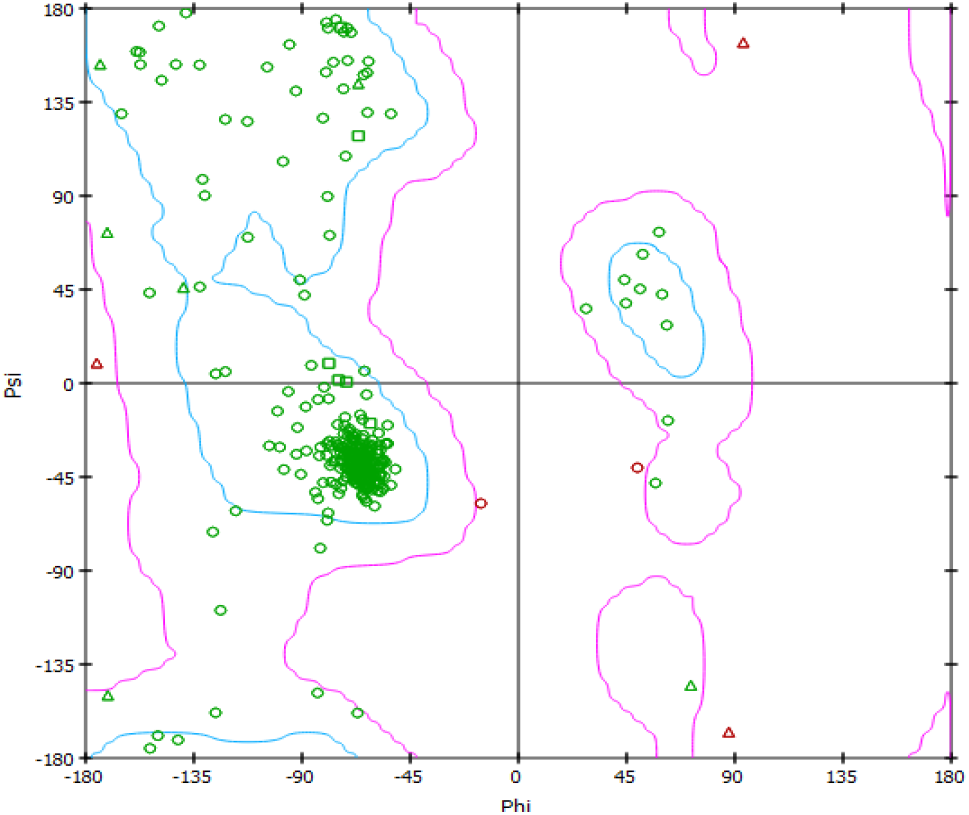
The Ramachandran plot for the APOE4 models after energy minimization.

### Preparation of target structure

APOE4 structure was prepared by adding hydrogen atoms were with hydrogen bond network optimization. Charges for standard residues were calculated using Amber 14SB force field and for ligands Gasteiger charges were used.(Pettersen et al., 2004).

### Selection of Ligands

#### Stabilizers

A study on NMR-based fragment screening and various other biophysical methods on APOE4 reported potential stabilizers (Petros et al., 2019), based on which, substructure search was performed. Similar structures containing stabilizers as their substructures were searched in the PubChem Database (https://pubchem.ncbi.nlm.nih.gov/) and identified total of 77 compounds. The structures of the eight stabilizers were also retrieved using ChemAxon Marvin Sketch (https://chemaxon.com/products/marvin).

#### Drugs

Various classes of drugs used in the treatment of CAD, including angiotensin-converting enzyme (ACE) inhibitors, beta blockers, bile acid sequestrates, cholesterol absorption inhibitors, factor Xa inhibitors, peripheral vasodilators, platelet aggregation inhibitors and statins were studied. The 3D conformers of the drugs were collected from the PubChem database and subjected for pre-docking processing.

#### Bioactive Molecules

Molecules from ChEMBL (https://www.ebi.ac.uk/chembl/) were also added to the list of compounds to test based on their bioactivity against APOE4 protein. The molecules retrieved were studied by western blot for inhibition of apoE4 expressed in Neuro2a cells and in rat PC12 cells assessed as reversal of mitochondrial impairment (Mahley & Huang, 2012).

#### Molecular docking simulations

Molecular docking is widely used for predicting the binding affinities for a number of ligands. In the present study, docking was performed with AutoDock Vina version 1.1.2 (Trott & Olson, 2009). All the ligand molecules were geometry optimized and energy minimized by Open Babel (http://openbabel.org) module using MMFF94 force field (O’Boyle et al., 2011) prior to docking. Binding site for docking, provided as a search volume, was derived from the co-crystallized stabilizer structures from PDB. Dimensions of the grid box were 28 × 26 × 28 with grid spacing of 1Å. Lowest energy conformations were chosen for further investigation. This procedure was applied to all the ligands and the selected conformations were analysed with receptor structure for interaction analysis. Docking for APOE3 as the receptor molecule was also performed using AutoDock Vina, with the grid dimension as 28 × 26 × 28 with grid spacing of 1 Å. The ligand conformation which showed the lowest docked energy (binding affinity) was chosen and compared with that of APOE4, as shown in Table 2.

**Table 2:**
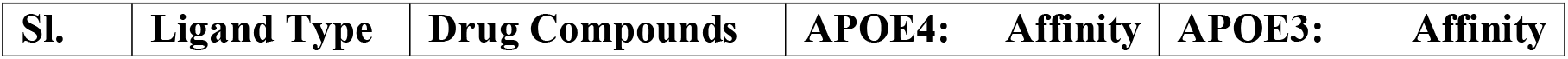
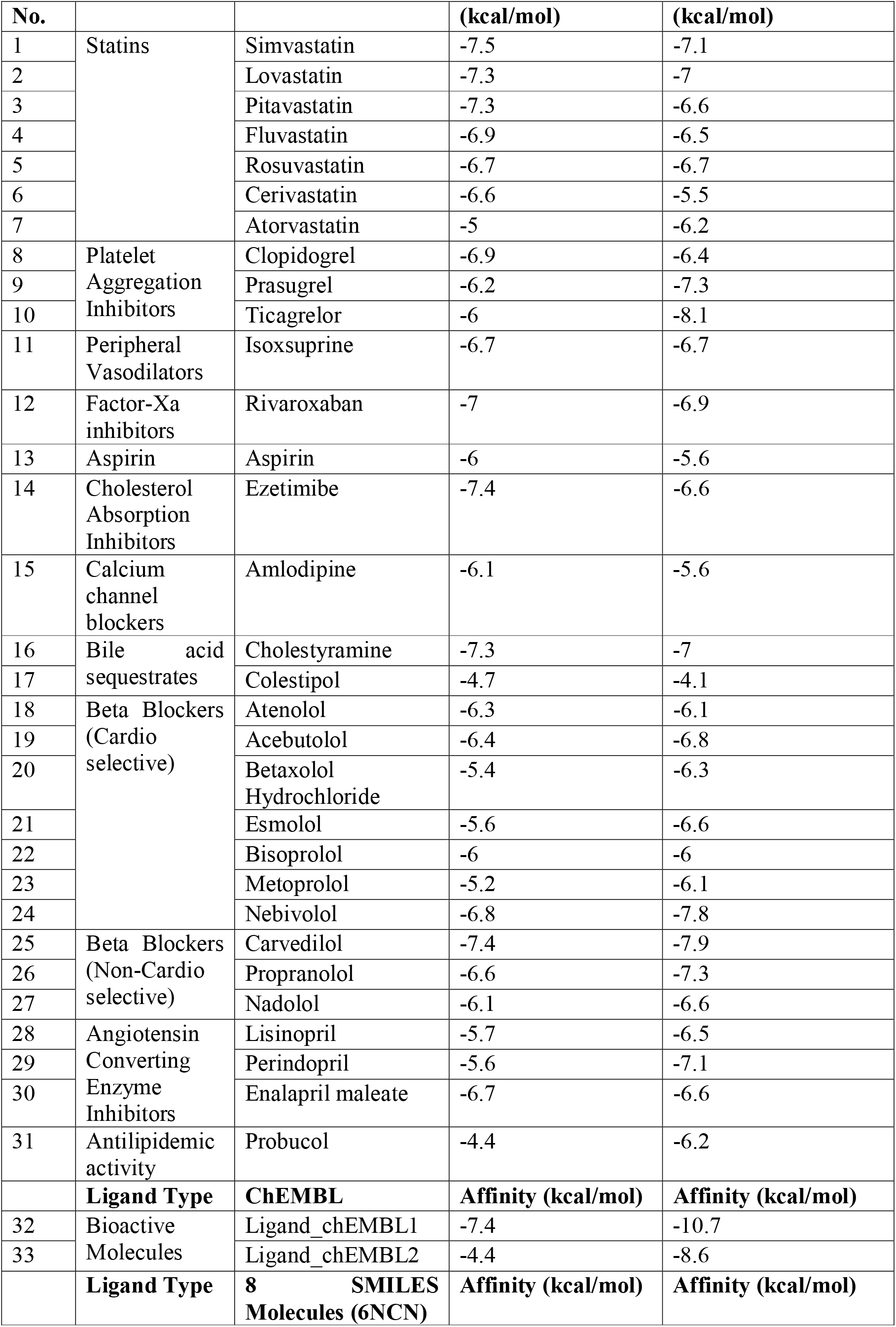
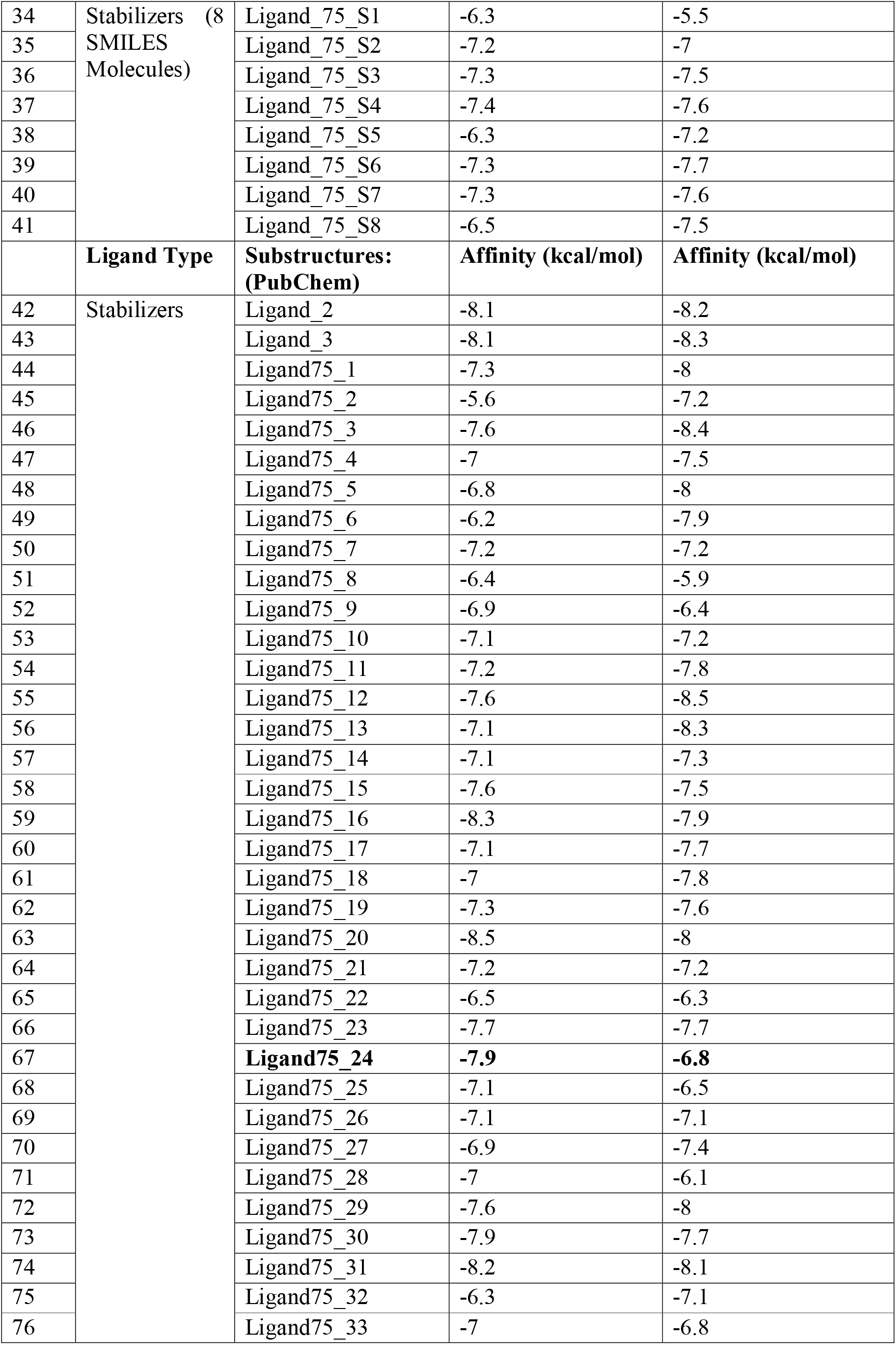
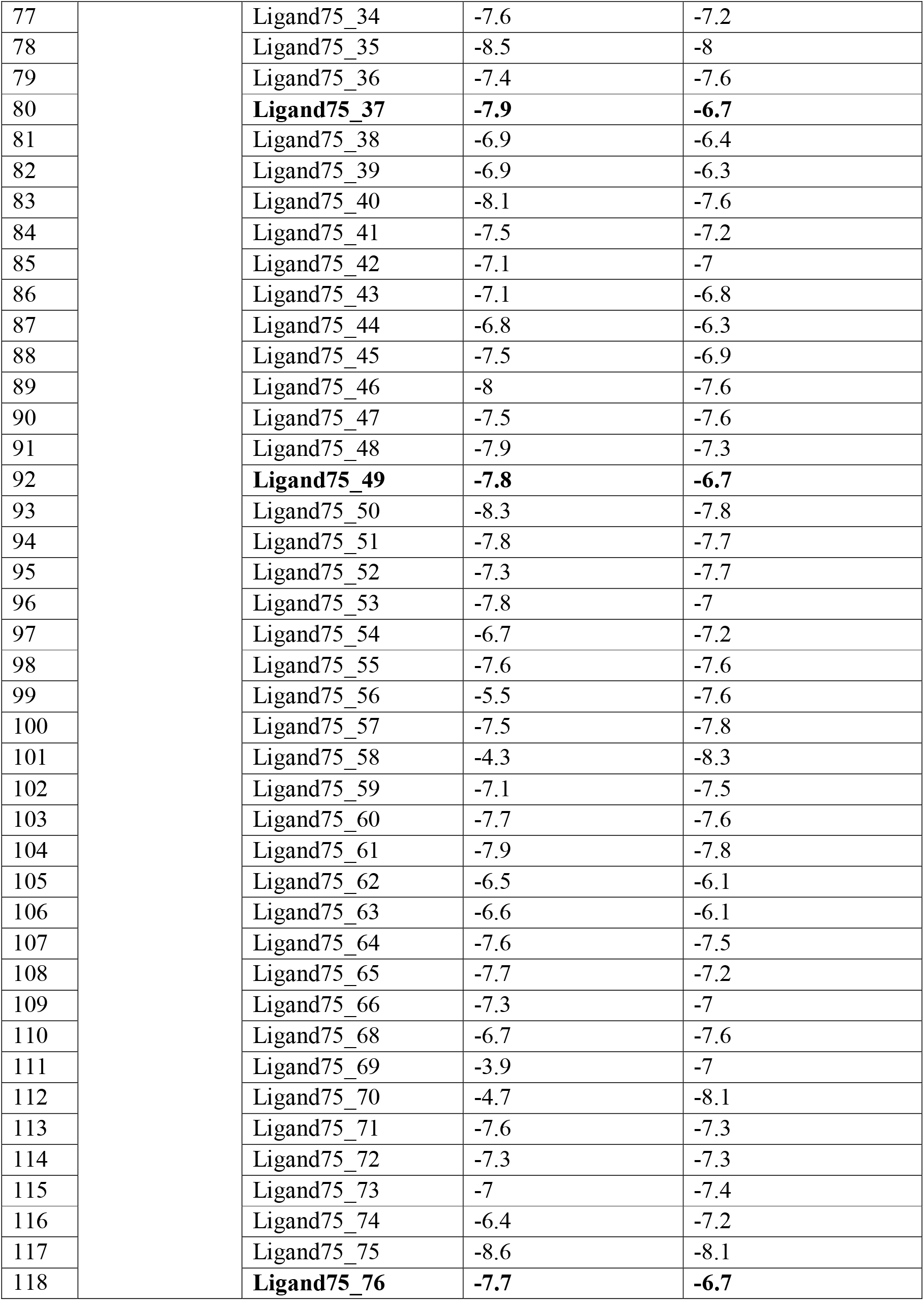
Differential binding affinities of APOE4 and APOE3 with various ligands.

| Sl. | Ligand Type | Drug Compounds | APOE4: Affinity | APOE3: Affinity |
| --- | --- | --- | --- | --- |

| No. |  |  | (kcal/mol) | (kcal/mol) |
| --- | --- | --- | --- | --- |
| 1 | Statins | Simvastatin | -7.5 | -7.1 |
| 2 |  | Lovastatin | -7.3 | -7 |
| 3 |  | Pitavastatin | -7.3 | -6.6 |
| 4 |  | Fluvastatin | -6.9 | -6.5 |
| 5 |  | Rosuvastatin | -6.7 | -6.7 |
| 6 |  | Cerivastatin | -6.6 | -5.5 |
| 7 |  | Atorvastatin | -5 | -6.2 |
| 8 | Platelet Aggregation Inhibitors | Clopidogrel | -6.9 | -6.4 |
| 9 |  | Prasugrel | -6.2 | -7.3 |
| 10 |  | Ticagrelor | -6 | -8.1 |
| 11 | Peripheral Vasodilators | Isoxsuprine | -6.7 | -6.7 |
| 12 | Factor-Xa inhibitors | Rivaroxaban | -7 | -6.9 |
| 13 | Aspirin | Aspirin | -6 | -5.6 |
| 14 | Cholesterol Absorption Inhibitors | Ezetimibe | -7.4 | -6.6 |
| 15 | Calcium channel blockers | Amlodipine | -6.1 | -5.6 |
| 16 | Bile acid sequestrates | Cholestyramine | -7.3 | -7 |
| 17 |  | Colestipol | -4.7 | -4.1 |
| 18 | Beta Blockers (Cardio selective) | Atenolol | -6.3 | -6.1 |
| 19 |  | Acebutolol | -6.4 | -6.8 |
| 20 |  | Betaxolol Hydrochloride | -5.4 | -6.3 |
| 21 |  | Esmolol | -5.6 | -6.6 |
| 22 |  | Bisoprolol | -6 | -6 |
| 23 |  | Metoprolol | -5.2 | -6.1 |
| 24 |  | Nebivolol | -6.8 | -7.8 |
| 25 | Beta Blockers (Non-Cardio selective) | Carvedilol | -7.4 | -7.9 |
| 26 |  | Propranolol | -6.6 | -7.3 |
| 27 |  | Nadolol | -6.1 | -6.6 |
| 28 | Angiotensin Converting Enzyme Inhibitors | Lisinopril | -5.7 | -6.5 |
| 29 |  | Perindopril | -5.6 | -7.1 |
| 30 |  | Enalapril maleate | -6.7 | -6.6 |
| 31 | Antilipidemic activity | Probucol | -4.4 | -6.2 |
|  | **Ligand Type** | **ChEMBL** | **Affinity (kcal/mol)** | **Affinity (kcal/mol)** |
| 32 | Bioactive Molecules | Ligand_chEMBL1 | -7.4 | -10.7 |
| 33 |  | Ligand_chEMBL2 | -4.4 | -8.6 |
|  | **Ligand Type** | **8 SMILES Molecules (6NCN)** | **Affinity (kcal/mol)** | **Affinity (kcal/mol)** |
| 34 | Stabilizers (8<br>SMILES<br>Molecules) | Ligand_75_S1 | -6.3 | -5.5 |
| 35 |  | Ligand_75_S2 | -7.2 | -7 |
| 36 |  | Ligand_75_S3 | -7.3 | -7.5 |
| 37 |  | Ligand_75_S4 | -7.4 | -7.6 |
| 38 |  | Ligand_75_S5 | -6.3 | -7.2 |
| 39 |  | Ligand_75_S6 | -7.3 | -7.7 |
| 40 |  | Ligand_75_S7 | -7.3 | -7.6 |
| 41 |  | Ligand_75_S8 | -6.5 | -7.5 |
|  | **Ligand Type** | **Substructures:**<br>**(PubChem)** | **Affinity (kcal/mol)** | **Affinity (kcal/mol)** |
| 42 | Stabilizers | Ligand_2 | -8.1 | -8.2 |
| 43 |  | Ligand_3 | -8.1 | -8.3 |
| 44 |  | Ligand75_1 | -7.3 | -8 |
| 45 |  | Ligand75_2 | -5.6 | -7.2 |
| 46 |  | Ligand75_3 | -7.6 | -8.4 |
| 47 |  | Ligand75_4 | -7 | -7.5 |
| 48 |  | Ligand75_5 | -6.8 | -8 |
| 49 |  | Ligand75_6 | -6.2 | -7.9 |
| 50 |  | Ligand75_7 | -7.2 | -7.2 |
| 51 |  | Ligand75_8 | -6.4 | -5.9 |
| 52 |  | Ligand75_9 | -6.9 | -6.4 |
| 53 |  | Ligand75_10 | -7.1 | -7.2 |
| 54 |  | Ligand75_11 | -7.2 | -7.8 |
| 55 |  | Ligand75_12 | -7.6 | -8.5 |
| 56 |  | Ligand75_13 | -7.1 | -8.3 |
| 57 |  | Ligand75_14 | -7.1 | -7.3 |
| 58 |  | Ligand75_15 | -7.6 | -7.5 |
| 59 |  | Ligand75_16 | -8.3 | -7.9 |
| 60 |  | Ligand75_17 | -7.1 | -7.7 |
| 61 |  | Ligand75_18 | -7 | -7.8 |
| 62 |  | Ligand75_19 | -7.3 | -7.6 |
| 63 |  | Ligand75_20 | -8.5 | -8 |
| 64 |  | Ligand75_21 | -7.2 | -7.2 |
| 65 |  | Ligand75_22 | -6.5 | -6.3 |
| 66 |  | Ligand75_23 | -7.7 | -7.7 |
| 67 |  | **Ligand75_24** | **-7.9** | **-6.8** |
| 68 |  | Ligand75_25 | -7.1 | -6.5 |
| 69 |  | Ligand75_26 | -7.1 | -7.1 |
| 70 |  | Ligand75_27 | -6.9 | -7.4 |
| 71 |  | Ligand75_28 | -7 | -6.1 |
| 72 |  | Ligand75_29 | -7.6 | -8 |
| 73 |  | Ligand75_30 | -7.9 | -7.7 |
| 74 |  | Ligand75_31 | -8.2 | -8.1 |
| 75 |  | Ligand75_32 | -6.3 | -7.1 |
| 76 |  | Ligand75_33 | -7 | -6.8 |

|  |  |  |  |
| --- | --- | --- | --- |
| 77 | Ligand75_34 | -7.6 | -7.2 |
| 78 | Ligand75_35 | -8.5 | -8 |
| 79 | Ligand75_36 | -7.4 | -7.6 |
| 80 | **Ligand75_37** | **-7.9** | **-6.7** |
| 81 | Ligand75_38 | -6.9 | -6.4 |
| 82 | Ligand75_39 | -6.9 | -6.3 |
| 83 | Ligand75_40 | -8.1 | -7.6 |
| 84 | Ligand75_41 | -7.5 | -7.2 |
| 85 | Ligand75_42 | -7.1 | -7 |
| 86 | Ligand75_43 | -7.1 | -6.8 |
| 87 | Ligand75_44 | -6.8 | -6.3 |
| 88 | Ligand75_45 | -7.5 | -6.9 |
| 89 | Ligand75_46 | -8 | -7.6 |
| 90 | Ligand75_47 | -7.5 | -7.6 |
| 91 | Ligand75_48 | -7.9 | -7.3 |
| 92 | **Ligand75_49** | **-7.8** | **-6.7** |
| 93 | Ligand75_50 | -8.3 | -7.8 |
| 94 | Ligand75_51 | -7.8 | -7.7 |
| 95 | Ligand75_52 | -7.3 | -7.7 |
| 96 | Ligand75_53 | -7.8 | -7 |
| 97 | Ligand75_54 | -6.7 | -7.2 |
| 98 | Ligand75_55 | -7.6 | -7.6 |
| 99 | Ligand75_56 | -5.5 | -7.6 |
| 100 | Ligand75_57 | -7.5 | -7.8 |
| 101 | Ligand75_58 | -4.3 | -8.3 |
| 102 | Ligand75_59 | -7.1 | -7.5 |
| 103 | Ligand75_60 | -7.7 | -7.6 |
| 104 | Ligand75_61 | -7.9 | -7.8 |
| 105 | Ligand75_62 | -6.5 | -6.1 |
| 106 | Ligand75_63 | -6.6 | -6.1 |
| 107 | Ligand75_64 | -7.6 | -7.5 |
| 108 | Ligand75_65 | -7.7 | -7.2 |
| 109 | Ligand75_66 | -7.3 | -7 |
| 110 | Ligand75_68 | -6.7 | -7.6 |
| 111 | Ligand75_69 | -3.9 | -7 |
| 112 | Ligand75_70 | -4.7 | -8.1 |
| 113 | Ligand75_71 | -7.6 | -7.3 |
| 114 | Ligand75_72 | -7.3 | -7.3 |
| 115 | Ligand75_73 | -7 | -7.4 |
| 116 | Ligand75_74 | -6.4 | -7.2 |
| 117 | Ligand75_75 | -8.6 | -8.1 |
| 118 | **Ligand75_76** | **-7.7** | **-6.7** |

## Results

Sequence alignment of target and template sequences performed by NCBI protein-protein BLAST analysis estimated the matches and similarity score. Among the accession IDs returned, the PDB IDs 2L7B (APOE3 of 307 amino acid) showed 98.33% percent identity and 94% query coverage and 1B68 (APOE4 of 191 amino acid) showed percentage identity of 99.48% and query coverage of 60% only. The full-length generated APOE4 sequence was compared with 2L7B and 1B68 by multiple sequence alignment tool and modelled by homology to generate a good quality model. Model scores derived from statistical potentials included z-DOPE score as −0.24; estimated root-mean-square deviation (RMSD) was 5.097 for the model. The structural assessment results for the 3D model showed 87.9% residues in most favoured region and only 0.7% residues in disallowed regions by Discovery Studio. The model validation was evaluated by Ramachandran Plot for the generated model as well as for the model which undergone energy minimization in presence of solvent as shown in Figure 3 a & b.

Results obtained on docking of 118 ligands with APOE4 compared to APOE3 showed different binding affinities, as shown in Table 2. Among various ligands, binding affinity of the substructure search for the SMILES: N=C(N)C1(CCC1)C2=CC(Cl)=CC=C2 showed better binding affinity towards the target protein. The highest binding affinity exhibited by Ligand75_75 for APOE4 was −8.6 kcal/mol. The dynamic structure of APOE3 undergoes domain interaction, but to a significantly lesser degree than APOE4 (Mahley & Huang, 2012) and hence differential binding affinities exhibited when docked with the two receptor molecules. Among all the compounds docked against the target protein, the ones which showed a differential binding affinity of 1 kcal/mol were selected for further investigations. Substructure viz, Ligand75_24 showed an affinity of −7.9 kcal/mol to APOE4 whereas that for APOE3 is −6.8 kcal/mol. The same is also observed with Ligand75_37 showing −7.9 kcal/mol with APOE4 and −6.7 kcal/mol for APOE3. Ligand75_49 shows affinity of −7.8 kcal/mol and −6.7 kcal/mol for APOE4 and APOE3 respectively whereas Ligand75_76 showed binding affinity of −7.7 kcal/mol and −6.7 kcal/mol for APOE4 and APOE3 respectively.

The receptor-ligand interactions were analysed using Discovery Studio showed the amino acid residues. Ligand75_24 showed hydrogen bonding with arginine 33 with glutamic acid at 37 and also involved in hydrophobic interactions (Figure 4). Glycine at 49 of the same ligands interacted through hydrogen bonding with glutamic acid at 45. The Ligand75_37 had its hydrogen bonding through arginine at 33 and Glycine at 49, whereas Tryptophan 52 and Leucine 167 showed hydrophobic interactions. Ligand75_49 had hydrogen bonding interactions from arginine at 33 to glycine at 49. Hydrophobic interactions for Ligand75_76 included tryptophan at 52 position and arginine at 33 position and leucine at 32, 48 and 167 position (Figure 4).

**Figure 4.a:**
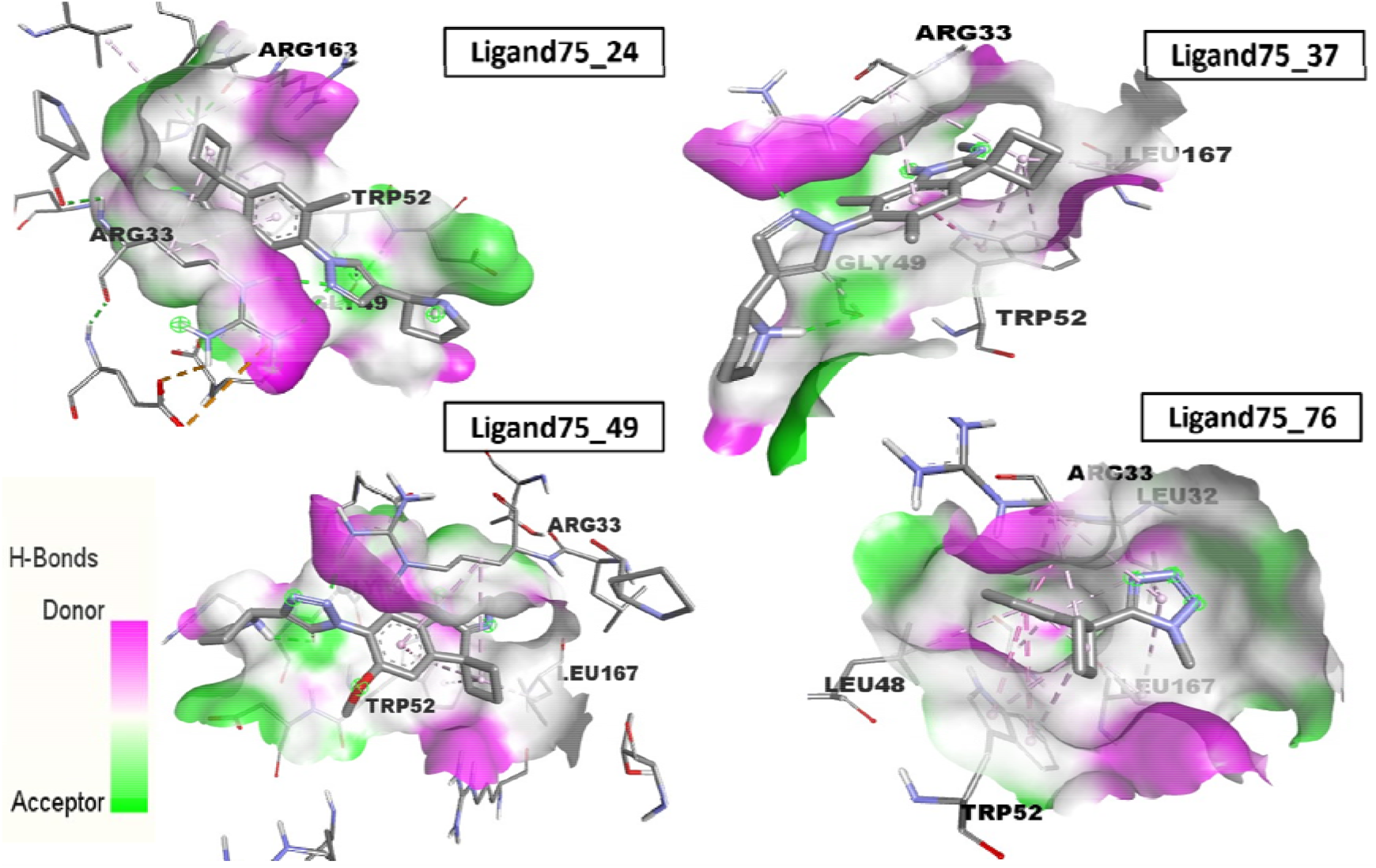
Receptor-ligand interactions visualized using discovery studio showing the amino acid residues interacting through hydrogen bonds with APOE4 receptor.

**Figure 4.b:**
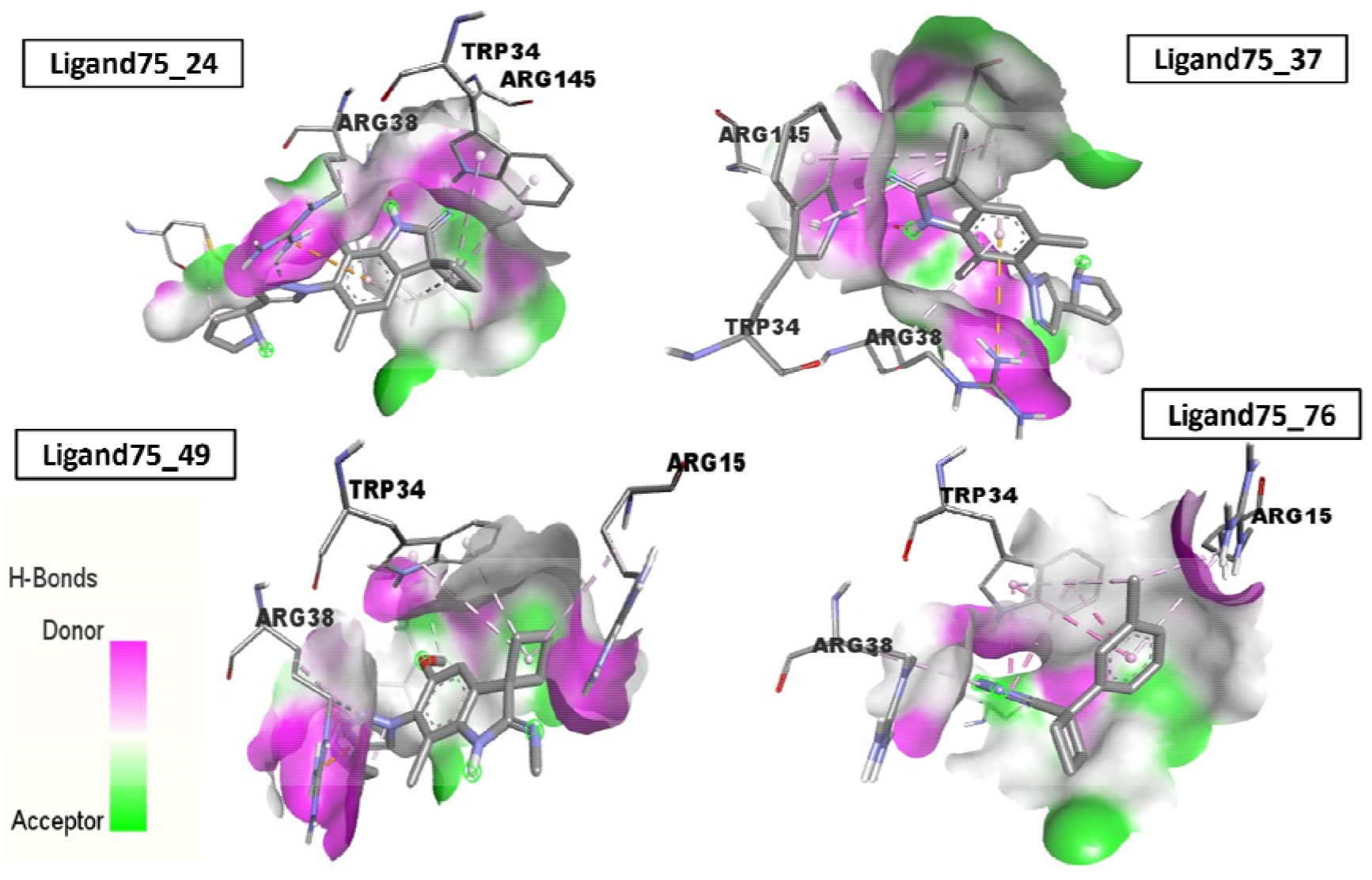
Receptor-ligand interactions visualized using discovery studio showing the amino acid residues interacting through hydrogen bonds with APOE3 receptor.

The most effective lipid-lowering drugs usually selected as the first-line therapy to lower total cholesterol (TC) and low density lipoprotein cholesterol (LDL-C) and increase the level of high density lipoprotein cholesterol (HDL-C) are the statins (M. Li et al., 2019) and widely prescribed for coronary heart disease. Simvastatin, a commonly used cholesterol lowering agent showed a binding affinity of −7.5 kcal/mol while Lovastatin, Pitavastatin and Cholestyramine showed an affinity of −7.3 kcal/mol. The well-known drug Rivaroxaban, is a direct inhibitor of the coagulation factor Xa with anticoagulant activity showed −7.0 kcal/mol; Ezetimibe, a cholesterol absorption inhibitor with lipid-lowering activity and Carvedilol, a nonselective beta-adrenoceptor blocking agent showed binding affinity of −7.4 kcal/mol (NCBI PubChem.). Interestingly, the affinity of the bioactive molecule viz, Ligand_chEMBL1 showed higher affinity for APOE3 (−10.7 kcal/mol) compared to APOE4 (−7.4 kcal/mol) in the present study, supporting that the ApoE3 and apoE4 bind equally well to the receptors, due to its positively charged arginine rich residues in the LDL receptor−binding region (Mahley & Huang, 2012).

Pharmacophore detection shared by selected input ligand molecules (Ligand75_24, Ligand75_37, Ligand75_49, Ligand75_76, Ezetimibe drug) was checked by PharmaGist (http://bioinfo3d.cs.tau.ac.il/PharmaGist/) which is a ligand-based method (Schneidman-Duhovny et al., 2008). Ezetimibe, a drug used in the treatment of CAD, shown to have significant improvement in lipid and lipoprotein profile (TC, LDL-C, triglycerides and HDL-C) in patients with APOE3 and APOE4 genotypes (Mark et al., 2007). Pharmacophore mapping of the selected ligands showed shared pharmacophore features between the 4 ligands and the Ezetimibe drug, containing 2 hydrophobic contacts and 2 hydrogen bond donors as shown in Figure 5.

**Figure 5:**
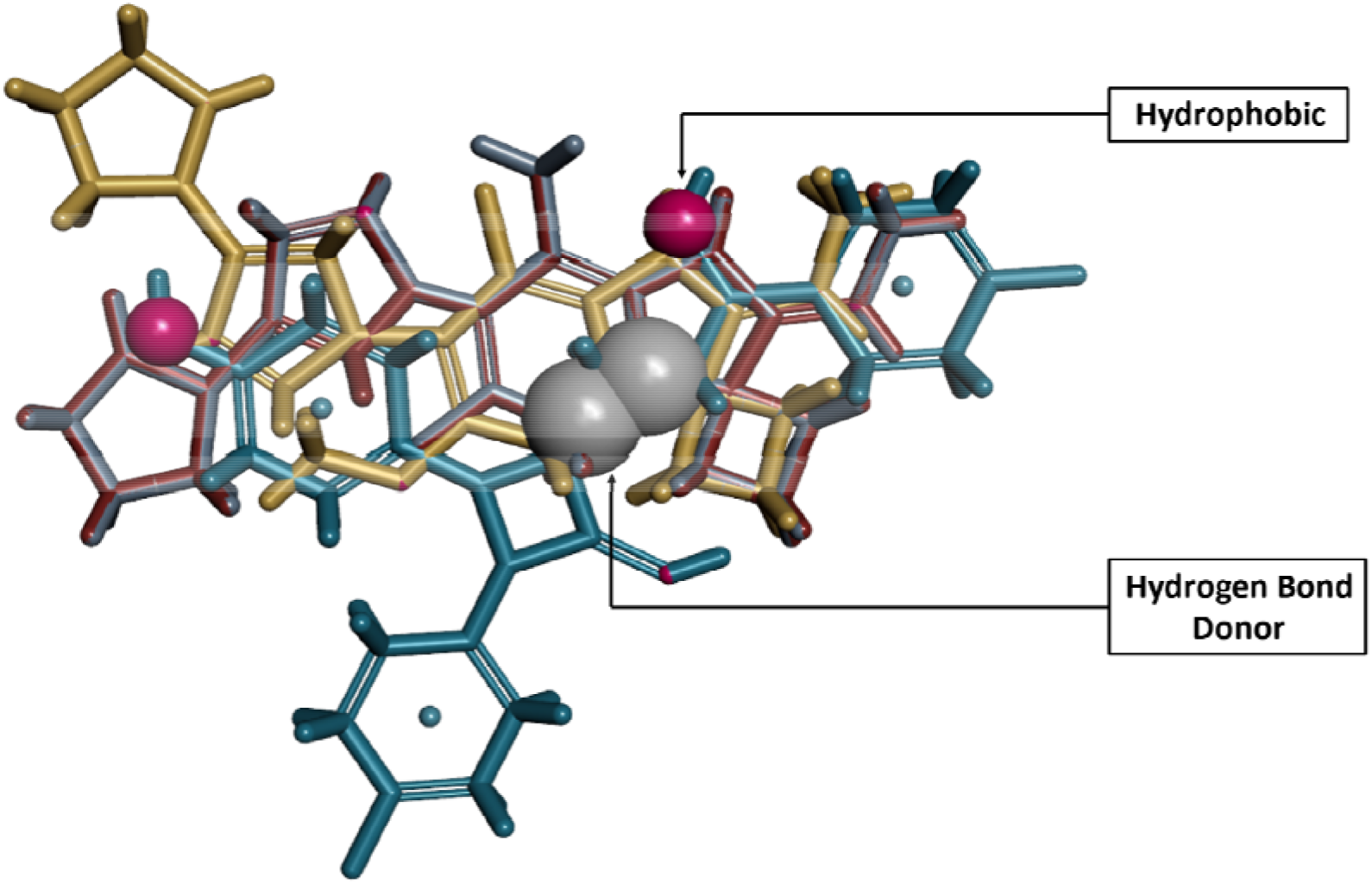
Pharmacophore mapping of the selected ligands viz, Ligand75_24, Ligand75_37, Ligand75_49, Ligand75_76 and Ezetimibe drug. Magenta spheres – Hydrophobic; Grey spheres – Hydrogen bond donor.

## Discussion

The aim of molecular docking is the precise prediction of the structure of ligand within the receptor binding site and correctly estimate the strength of binding. To investigate the effective drugs for CAD, various compounds and substructures against the targets for CAD was studied utilizing molecular docking. It is well known that there is no exact treatment to prevent or cure CAD, currently, there are several approved drugs alone or in combination that have been used to reduce symptoms and briefly improve the condition. The present study aimed at evaluating all the existing drugs molecules and compounds as ligands against the APOE4, a potential predictor for CAD. Among the drugs, Ezetimibe has been earlier studied and found to be used in the treatment of hypercholesterolemia (Burnett & Huff, 2006) and is effective for lowering LDL-C and non HDL-C as monotherapy or in combination with statins (Al-Shaer et al., 2004). In the present study, the drug had a binding affinity of −7.4 kcal/mol to APOE4, presenting its candidature. Effects of the combination of drugs like rivaroxaban and aspirin were also tested in patients with stable CAD or peripheral artery disease (PAD) (Kruger et al., 2018). Rivaroxaban reported with −7.0 kcal/mol affinity to APOE4 in the present study and the drug is under a recent randomized clinical trial among patients with atrial fibrillation and stable coronary artery disease (Bavry & Bhatt, 2020).

The differential binding energies exhibited from the docking indicated side chain stabilization or backbone dynamics that included the interactions mediated by hydrogen bonds and hydrophobic bonds. Among all the ligands studied, four substructures of ligands named as Ligand75 showed the most favourable binding affinity towards APOE4. The receptor and ligand interaction exhibited mostly arginine residues for hydrogen bonding; hydrogen bonding being reported to promote high-affinity receptor-ligand interactions, a principle important for drug design research (Chen et al., 2016). Our study found arginine, a positively charged amino acid involved in salt-bridge formation with negatively charged glutamic acid creating stabilising hydrogen bonds. This indicates the probable activity of the amino acid for the protein to bind to LDL-receptor. Studies suggested that the preferential binding of APOE4 to VLDL and APOE3 to HDL depends on the molecular basis of APOE isoform which rests in region from 261–272, involved in lipid binding. Residues in peptides 15–30, 116–123, and 271–279 show greater exchange in APOE4 relative to APOE3, suggesting greater dynamic motion (Frieden & Garai, 2012).

The interactions between the ligand and target protein could predict the ligand binding-conformation through differential binding affinities, which is influenced by non-covalent intermolecular interactions such as hydrogen bonding, electrostatic interactions, hydrophobic and Van der Waals forces between the two molecules(Pantsar & Poso, 2018; Saxena et al., 2009) and this is a key to understand the driving biological processes, structural biology, and structure-function relationships (MalvernPanalytical). The hydrogen bonding and ionic terms are both dependent on the geometry of the interaction, with large deviations from ideal geometries being penalised. The binding affinities calculated for studied Ligands docked to the 3D structures of APOE4 and APOE3 showed 1 kcal/mol difference in binding between these receptor variants, reported in Table 2. A change of around 1.4 kcal/mol in the free energy corresponds to a tenfold change in the free energy of binding (Leach & Gillet, 2007). Pharmacophore detection, fundamental to understand molecular interactions, serves as an ideal layout in novel drug design studies (Schneidman-Duhovny et al., 2008). The selected ligands in the current investigation indicated common features distributed in the 3D space suggesting Ligand75 modulators hold significant interactions and can be candidate leads for drug development.

Holloway et al. explained the pharmacodynamics of the drug-receptor properties as distinctive that indicates saturability, selectivity and the binding affinity. Saturability as the concentration of drug molecules occupying the maximum number of binding sites in the receptor. The degree to which a receptor binds with a specific drug is called receptor selectivity that depends on the receptor and on the size, shape, and bioelectrical charge of the drug. The strength of the interaction between the receptor and the drug molecule is the binding affinity, that occur by intermolecular forces, such as ionic bonds, hydrogen bonds and Van der Waals forces and this has a fundamental role in drug development. High-affinity ligand binding results indicate that a receptor require a lower drug concentration for full saturation. This can be understood from the concept of agonists and antagonists that could bind to the same receptor but differ in their affinity as high affinity agonist and low affinity antagonist could lead to an overwhelming drug effect (Holloway & Peirce, 1998). Ligands with weaker binding affinities can be characterized by their high dissociation rates and transient interactions with the target molecule (Wang et al., 2016).

The present study reported binding affinities of various classes of ligands to APOE4 and APOE3 predicted by the docking which suggests arginine residue with hydrogen bonding and hydrophobic interactions might affect the LDL receptor binding site of the APOE4, indicating distinctive lipid and lipoprotein binding activities. Although binding affinity data alone does not determine the overall potency of a drug but in the present study the selected ligands identified does reflect a candidature for its efficacy, which needs further investigations on the molecular dynamics. It can also be said that the altered structural conformation of APOE4 influenced by intramolecular domain interaction contributes to differential binding affinities and hence, a probable answer to APOE4 as a contributing factor towards CAD pathogenesis.

## Conclusion

Molecular modelling and docking are a promising aspect of drug discovery given the time frame required to investigate possible therapeutics in CAD. The present study has predicted the protein–ligand binding affinities indicating that receptor binding abilities of APOE4 can lead to have insights on structural conformity of APOE4 and its correlated functional aspects. Docking studies revealed the mode of binding and the binding energy data presented a good picture of ligands’ affinity and fitting inside the binding pocket of APOE4. However, more investigations on these bindings are required to improve and understand drug-receptor interactions. The selected Ligand75 modulators could be potential leads to evolve as candidate drug candidates against APOE4 in the treatment of atherosclerosis in CAD. Further analysis of the variation of the docked protein structure, molecular dynamic simulation can be performed to generate a dynamic structure for binding analysis.

## Supplementary Material

Table attached.

## Disclosure / Funding

This research received no specific grant from any funding agency in the public, commercial, or not-for-profit sectors.

## Conflicts of interest

The authors declare no conflict of interest.

## Notes

### Competing Interest Statement

The authors have declared no competing interest.

## References

Afroze, D., Yousuf, A., Tramboo, N. A., Shah, Z. A., & Ahmad, A. (2016). ApoE gene polymorphism and its relationship with coronary artery disease in ethnic Kashmiri population. Clinical and Experimental Medicine, 16(4), 551–556. https://doi.org/10.1007/s10238-015-0389-7

Al-Shaer, M. H., Choueiri, N. E., & Suleiman, E. S. (2004). The pivotal role of cholesterol absorption inhibitors in the management of dyslipidemia. In Lipids in Health and Disease (Vol. 3, Issue 1, p. 22). BioMed Central. https://doi.org/10.1186/1476-511X-3-22

Bavry, A. A., & Bhatt, D. L. (n.d.). Atrial Fibrillation and Ischemic Events With Rivaroxaban in Patients With Stable Coronary Artery Disease - American College of Cardiology. 2020. Retrieved December 29, 2020, from https://www.acc.org/latest-in-cardiology/clinical-trials/2019/09/01/11/05/afire

Boulenouar, H., Benchekor, S. M., Meroufel, D. N., Hetraf, S. A. L., Djellouli, H. O., Hermant, X., Grenier-Boley, B., Medjaoui, I. H., Mehtar, N. S., Amouyel, P., Houti, L., Meirhaeghe, A., & Goumidi, L. (2013). Impact of APOE gene polymorphisms on the lipid profile in an Algerian population. Lipids in Health and Disease, 12(1). https://doi.org/10.1186/1476-511X-12-155

Burnett, J. R., & Huff, M. W. (2006). Cholesterol absorption inhibitors as a therapeutic option for hypercholesterolaemia. In Expert Opinion on Investigational Drugs (Vol. 15, Issue 11, pp. 1337–1351). Expert Opin Investig Drugs. https://doi.org/10.1517/13543784.15.11.1337

CDC. (n.d.). Coronary Artery Disease | cdc.gov. Retrieved November 30, 2020, from https://www.cdc.gov/heartdisease/coronary_ad.htm

Chen, C. Y.-C. (2013). A Novel Integrated Framework and Improved Methodology of Computer-Aided Drug Design. Current Topics in Medicinal Chemistry, 13(9), 965–988. https://doi.org/10.2174/1568026611313090002

Chen, D., Oezguen, N., Urvil, P., Ferguson, C., Dann, S. M., & Savidge, T. C. (2016). Regulation of protein-ligand binding affinity by hydrogen bond pairing. Science Advances, 2(3), e1501240. https://doi.org/10.1126/sciadv.1501240

Chou, C. Y., Jen, W. P., Hsieh, Y. H., Shiao, M. S., & Chang, G. G. (2006). Structural and functional variations in human apolipoprotein E3 and E4. Journal of Biological Chemistry, 281(19), 13333–13344. https://doi.org/10.1074/jbc.M511077200

Coronary Heart Disease | NHLBI, NIH. (n.d.). Retrieved November 30, 2020, from https://www.nhlbi.nih.gov/health-topics/coronary-heart-disease

Freeman, M. W. (2006). Lipid metabolism and coronary artery disease. Principles of Molecular Medicine, 130–137. https://doi.org/10.1007/978-1-59259-963-9_15

Frieden, C., & Garai, K. (2012). Structural differences between apoE3 and apoE4 may be useful in developing therapeutic agents for Alzheimer’s disease. Proceedings of the National Academy of Sciences of the United States of America, 109(23), 8913–8918. https://doi.org/10.1073/pnas.1207022109

Frieden, C., Wang, H., & Ho, C. M. W. (2017). A mechanism for lipid binding to apoE and the role of intrinsically disordered regions coupled to domain-domain interactions. Proceedings of the National Academy of Sciences of the United States of America, 114(24), 6292–6297. https://doi.org/10.1073/pnas.1705080114

Gopalakrishnan, A., Sivadasanpillai, H., Ganapathi, S., Mohanan Nair, K., Sivasubramonian, S., & Valaparambil, A. (2020). Clinical profile & long-term natural history of symptomatic coronary artery disease in young patients (30 yr). Indian Journal of Medical Research, 152(3), 263. https://doi.org/10.4103/ijmr.IJMR_1090_18

Holloway, F. A., & Peirce, J. M. (1998). Fundamental Psychopharmacology. In Comprehensive Clinical Psychology (pp. 173–206). Elsevier. https://doi.org/10.1016/b0080-4270(73)00176-0

Krieger, E., Joo, K., Lee, J., Lee, J., Raman, S., Thompson, J., Tyka, M., Baker, D., & Karplus, K. (2009). Improving physical realism, stereochemistry, and side-chain accuracy in homology modeling: Four approaches that performed well in CASP8. In Proteins: Structure, Function and Bioinformatics (Vol. 77, Issue SUPPL. 9, pp. 114–122). https://doi.org/10.1002/prot.22570

Kruger, P. C., Eikelboom, J. W., & Yusuf, S. (2018). Rivaroxaban with or without aspirin for prevention of cardiovascular disease. In Coronary Artery Disease (Vol. 29, Issue 5, pp. 361–365). Lippincott Williams and Wilkins. https://doi.org/10.1097/MCA.0000000000000605

Lamia, L. F., Sharif, F. A., & Abed, A. A. (2011). Relationship between ApoE gene polymorphism and coronary heart disease in Gaza Strip. Journal of Cardiovascular Disease Research, 2(1), 29–35. https://doi.org/10.4103/0975-3583.78584

Leach, A. R., & Gillet, V. J. (2007). An introduction to chemoinformatics. In An Introduction To Chemoinformatics. https://doi.org/10.1007/978-1-4020-6291-9

Li, H., Dhanasekaran, P., Alexander, E. T., Rader, D. J., Phillips, M. C., & Lund-Katz, S. (2013). Molecular mechanisms responsible for the differential effects of ApoE3 and ApoE4 on plasma lipoprotein-cholesterol levels. Arteriosclerosis, Thrombosis, and Vascular Biology, 33(4), 687–693. https://doi.org/10.1161/ATVBAHA.112.301193

Li, M., Wang, X., Li, X., Chen, H., Hu, Y., Zhang, X., Tang, X., Miao, Y., Tian, G., & Shang, H. (2019). Statins for the Primary Prevention of Coronary Heart Disease. BioMed Research International, 2019. https://doi.org/10.1155/2019/4870350

Lionta, E., Spyrou, G., Vassilatis, D., & Cournia, Z. (2014). Structure-Based Virtual Screening for Drug Discovery: Principles, Applications and Recent Advances. Current Topics in Medicinal Chemistry, 14(16), 1923–1938. https://doi.org/10.2174/1568026614666140929124445

Mahley, R. W. (2016). Apolipoprotein E: from cardiovascular disease to neurodegenerative disorders. Journal of Molecular Medicine, 94(7), 739–746. https://doi.org/10.1007/s00109-016-1427-y

Mahley, R. W., & Huang, Y. (2012). Small-Molecule structure correctors target abnormal protein structure and function: Structure corrector rescue of apolipoprotein E4-associated neuropathology. In Journal of Medicinal Chemistry (Vol. 55, Issue 21, pp. 8997–9008). J Med Chem. https://doi.org/10.1021/jm3008618

Mahley, R. W., & Rall, S. C. (2000). Apolipoprotein E: Far more than a lipid transport protein. Annual Review of Genomics and Human Genetics, 1(2000), 507–537. https://doi.org/10.1146/annurev.genom.1.1.507

MalvernPanalytical. (n.d.). Binding Affinity | Dissociation Constant | Malvern Panalytical. Retrieved January 7, 2021, from https://www.malvernpanalytical.com/en/products/measurement-type/binding-affinity

Mark, L., Dani, G., Fazekas, Ö., Szüle, O., Kovacs, H., & Katona, A. (2007). Effects of ezetimibe on lipids and lipoproteins in patients with hypercholesterolemia and different apolipoprotein E genotypes. Current Medical Research and Opinion, 23(7), 1541–1548. https://doi.org/10.1185/030079907X199817

Moriarty, P. M. (2009). Association of ApoE and HDL-C with cardiovascular and cerebrovascular disease: Potential benefits of LDL-apheresis therapy. Future Lipidology, 4(3), 311–329. https://doi.org/10.2217/CLP.09.21

NCBI PubChem. (n.d.). PubChem. Retrieved January 7, 2021, from https://pubchem.ncbi.nlm.nih.gov/

Ohashi, R., Mu, H., Wang, X., Yao, Q., & Chen, C. (2005). Reverse cholesterol transport and cholesterol efflux in atherosclerosis. QJM - Monthly Journal of the Association of Physicians, 98(12), 845–856. https://doi.org/10.1093/qjmed/hci136

Pantsar, T., & Poso, A. (2018). Binding affinity via docking: Fact and fiction. In Molecules (Vol. 23, Issue 8, p. 1DUMMY). MDPI AG. https://doi.org/10.3390/molecules23081899

Petros, A. M., Korepanova, A., Jakob, C. G., Qiu, W., Panchal, S. C., Wang, J., Dietrich, J. D., Brewer, J. T., Pohlki, F., Kling, A., Wilcox, K., Lakics, V., Bahnassawy, L., Reinhardt, P., Partha, S. K., Bodelle, P. M., Lake, M., Charych, E. I., Stoll, V. S., ... Mohler, E. G. (2019). Fragment-Based Discovery of an Apolipoprotein E4 (apoE4) Stabilizer. Journal of Medicinal Chemistry, 62(8), 4120–4130. https://doi.org/10.1021/acs.jmedchem.9b00178

Pettersen, E. F., Goddard, T. D., Huang, C. C., Couch, G. S., Greenblatt, D. M., Meng, E. C., & Ferrin, T. E. (2004). UCSF Chimera - A visualization system for exploratory research and analysis. Journal of Computational Chemistry, 25(13), 1605–1612. https://doi.org/10.1002/jcc.20084

Phillips, M. C. (2014). Apolipoprotein e isoforms and lipoprotein metabolism. IUBMB Life, 66(9), 616–623. https://doi.org/10.1002/iub.1314

Prakash, N., & Devangi, P. (2010). Drug Discovery. Journal of Antivirals and Antiretrovirals, 2(4), 063–068. https://doi.org/10.4172/jaa.1000025

Rall, S. C., Weisgraber, K. H., & Mahley, W. (1982). Human E. Biological Chemistry, 257(8).

Ranjith, N., Pegoraro, R., Rom, L., Rajput, M., & Naidoo, D. (2004). Sabinet | Lp(a) and apoE polymorphisms in young South African Indians with myocardial infarctionl_: cardiovascular topics. Cardiovascular Journal of South Africa. https://journals.co.za/content/cardio/15/3/EJC23918

Rezeli, M., Zetterberg, H., Blennow, K., Brinkmalm, A., Laurell, T., Hansson, O., & Marko-Varga, G. (2015). Quantification of total apolipoprotein E and its specific isoforms in cerebrospinal fluid and blood in Alzheimer’s disease and other neurodegenerative diseases. EuPA Open Proteomics, 8, 137–143. https://doi.org/10.1016/j.euprot.2015.07.012

Safieh, M., Korczyn, A. D., & Michaelson, D. M. (2019). ApoE4: an emerging therapeutic target for Alzheimer’s disease. BMC Medicine, 17(1), 1–17. https://doi.org/10.1186/s12916-019-1299-4

Saxena, A., Wong, D., Diraviyam, K., & Sept, D. (2009). The Basic Concepts of Molecular Modeling. In Methods in Enzymology (1st ed., Vol. 467, Issue C). Elsevier Inc. https://doi.org/10.1016/S0076-6879(09)67012-9

Schneidman-Duhovny, D., Dror, O., Inbar, Y., Nussinov, R., & Wolfson, H. J. (2008). PharmaGist: a webserver for ligand-based pharmacophore detection. Nucleic Acids Research, 36(Web Server issue), 223–228. https://doi.org/10.1093/nar/gkn187

Singh, P. P., Singh, M., & Mastana, S. S. (2006). APOE distribution in world populations with new data from India and the UK. Annals of Human Biology, 33(3), 279–308. https://doi.org/10.1080/03014460600594513

Singh, P., Singh, M., Bhatnagar, D., Kaur, T., & Gaur, S. (2008). Apolipoprotein E polymorphism and its relation to plasma lipids in coronary heart disease. Indian Journal of Medical Sciences, 62(3), 105–112. https://doi.org/10.4103/0019-5359.39613

Trott, O., & Olson, A. J. (2009). AutoDock Vina: Improving the speed and accuracy of docking with a new scoring function, efficient optimization, and multithreading. Journal of Computational Chemistry, NA-NA. https://doi.org/10.1002/jcc.21334

Wang, J., Guo, Z., Fu, Y., Wu, Z., Huang, C., Zheng, C., Shar, P. A., Wang, Z., Xiao, W., & Wang, Y. (2016). Weak-binding molecules are not drugs?—toward a systematic strategy for finding effective weak-binding drugs. Briefings in Bioinformatics, 18(2), bbw018. https://doi.org/10.1093/bib/bbw018

Weisgraber, K. H. (1994). Apolipoprotein E: Structure-function relationships. Advances in Protein Chemistry, 45, 249–302. https://doi.org/10.1016/s0065-3233(08)60642-7

