## Supplementary Table for "Identification of APOE4 modulators, targeted therapeutic candidates in Coronary Artery Disease, using Molecular Docking studies"

| Ligand75_1.sdf | Clc1ccc(C2(c3nnc4n3CCCCCC4)CCC2)cc1Cl |
| --- | --- |
| Ligand75_10.sdf | C/N=C1\Nc2cc(C#CCN3CCCCC3)c(Cl)cc2C12CCC2 |
| Ligand75_11.sdf | O=C(Nc1ccc(C2(c3nc4ncc(Br)cc4[nH]3)CCC2)cc1Cl)c1cccc(Br)c1 |
| Ligand75_12.sdf | N#Cc1cccc(C(=O)Nc2ccc(C3(c4nc5ncc(C#N)cc5[nH]4)CCC3)cc2Cl)c1 |
| Ligand75_13.sdf | CC(C)(O)c1ccc(-c2cc(Cl)cc(C3(C(=N)N)CCC3)c2)cc1 |
| Ligand75_14.sdf | CN(Cc1nc(C2(c3cccc(Cl)c3)CCC2)no1)S(C)(=O)=O |
| Ligand75_15.sdf | CCC(=O)N1CCCCC1c1nc(C2(c3cccc(Cl)c3)CCC2)no1 |
| Ligand75_16.sdf | CCC(=O)N1CCCC(c2nc(C3(c4cccc(Cl)c4)CCC3)no2)C1 |
| Ligand75_17.sdf | O=C1CCCN1CCc1nc(C2(c3cccc(Cl)c3)CCC2)no1 |
| Ligand75_18.sdf | CC(CS(C)(=O)=O)c1nc(C2(c3cccc(Cl)c3)CCC2)no1 |
| Ligand75_19.sdf | CC(C)(c1nc(C2(c3cccc(Cl)c3)CCC2)no1)S(C)(=O)=O |
| Ligand75_2.sdf | FC(F)(F)c1cccc(CN2CCN(CCNCc3nc(C4(c5cccc(Cl)c5)CCC4)no3)CC2)c1 |
| Ligand75_20.sdf | C/N=C1\Nc2cc(-n3cc([C@H]4CCCN4)nn3)c(Cl)cc2C12CCC2 |
| Ligand75_21.sdf | C/N=C1\Nc2cc(-c3cc4n(n3)CCNC4)c(Cl)cc2C12CCC2 |
| Ligand75_22.sdf | N=C(N)C1(c2cc(Cl)cc(Br)c2)CCC1 |
| Ligand75_23.sdf | NC(=O)c1nc(C2(c3cccc(Cl)c3)CCC2)no1 |
| Ligand75_24.sdf | C/N=C1\Nc2c(cc(Cl)c(-n3cc(C4CCCN4)cn3)c2C)C12CCC2 |
| Ligand75_25.sdf | C/N=C1\Nc2c(cc(Cl)c(-n3cc(CNC)cn3)c2Cl)C12CCC2 |
| Ligand75_26.sdf | C/N=C1\Nc2cc(N(N)/C=C(\N)C3CCCCN3)c(Cl)cc2C12CCC2 |
| Ligand75_27.sdf | C/N=C1\Nc2cc(N(N)/C=C(\N)C3CCCN3C)c(Cl)cc2C12CCC2 |
| Ligand75_28.sdf | C/N=C1\Nc2c(cc(C)c(C)c2Cl)C12CCC2 |
| Ligand75_29.sdf | C/N=C1\Nc2cc(-n3ncc4c3CCN(C)C4)c(Cl)cc2C12CCC2 |
| Ligand75_3.sdf | FC(F)(F)c1cccc(C2CCN(CCNCc3nc(C4(c5cccc(Cl)c5)CCC4)no3)CC2)c1 |
| Ligand75_30.sdf | CC1(c2nc(C3(c4cccc(Cl)c4)CCC3)no2)CCNC(=O)C1 |
| Ligand75_31.sdf | Cc1cc(Cc2nc(C3(c4cccc(Cl)c4)CCC3)no2)no1 |
| Ligand75_32.sdf | C/N=C1\Nc2cc(-n3cc(CN(C)C)cn3)c(Cl)cc2C12CCC2 |
| Ligand75_33.sdf | C/N=C1\Nc2c(cc(Cl)c(-n3cc(C4CCCN4)cn3)c2Cl)C12CCC2 |
| Ligand75_34.sdf | C/N=C1\Nc2c(cc(Cl)c(-n3cc([C@H]4CCCN4)cn3)c2Cl)C12CCC2 |
| Ligand75_35.sdf | C/N=C1\Nc2cc(-n3cc([C@H]4CCCN4)cn3)c(Cl)cc2C12CCC2 |
| Ligand75_36.sdf | C/N=C1\Nc2cc(-n3cc([C@@H]4CCCN4)cn3)c(Cl)cc2C12CCC2 |
| Ligand75_37.sdf | C/N=C1\Nc2c(cc(Cl)c(-n3cc([C@@H]4CCCN4)cn3)c2Cl)C12CCC2 |
| Ligand75_38.sdf | C/N=C1\Nc2c(cc(Cl)c(-n3cccn3)c2Cl)C12CCC2 |
| Ligand75_39.sdf | C/N=C1\Nc2c(cc(OC)c(-n3cc(CNC)cn3)c2Cl)C12CCC2 |
| Ligand75_4.sdf | COc1ccccc1C1CCN(CCNCc2nc(C3(c4ccc(Cl)c(Cl)c4)CCC3)no2)CC1 |
| Ligand75_40.sdf | C/N=C1\Nc2cc(-n3cc(C4CCCN4)cn3)c(Cl)cc2C12CCC2 |
| Ligand75_41.sdf | C/N=C1\Nc2cc(-n3cc(CNC)cn3)c(Cl)cc2C12CCC2 |
| Ligand75_42.sdf | C/N=C1\Nc2c(cc(Cl)c(-n3cc(CNC)cn3)c2F)C12CCC2 |
| Ligand75_43.sdf | C/N=C1\Nc2c(cc(Cl)c(N3CC([C@@H]4CCCN4)C=N3)c2Cl)C12CCC2 |
| Ligand75_44.sdf | C/N=C1\Nc2c(cc(Cl)c(-n3cc(C)cn3)c2Cl)C12CCC2 |
| Ligand75_45.sdf | C/N=C1\Nc2cc(-n3cccn3)c(Cl)cc2C12CCC2 |
| Ligand75_46.sdf | C/N=C1\Nc2cc(-n3cc(C4CCCN4)nn3)c(Cl)cc2C12CCC2 |
| Ligand75_47.sdf | C/N=C1\Nc2cc(-n3cc([C@@H]4CCCN4)nn3)c(Cl)cc2C12CCC2 |
| Ligand75_48.sdf | C/N=C1\Nc2cc(-n3cc4c(n3)CCCNC4)c(Cl)cc2C12CCC2 |
| Ligand75_49.sdf | C/N=C1\Nc2c(cc(OC)c(-n3cc([C@@H]4CCCN4)nn3)c2Cl)C12CCC2 |
| Ligand75_5.sdf | FC(F)(F)c1cccc(C2CCN(CCNCc3nc(C4(c5ccc(Cl)c(Cl)c5)CCC4)no3)CC2)c1 |
| Ligand75_50.sdf | C/N=C1\Nc2cc(-n3cc([C@@H]4CCCCN4)nn3)c(Cl)cc2C12CCC2 |
| Ligand75_51.sdf | C/N=C1\Nc2cc(-n3cc(C4CCCCN4)nn3)c(Cl)cc2C12CCC2 |
| Ligand75_52.sdf | C/N=C1\Nc2cc(-n3ncc4c3CCNC4)c(Cl)cc2C12CCC2 |
| Ligand75_53.sdf | C/N=C1\Nc2c(cc(Cl)c(-n3cc(C4CCCC4)cn3)c2Cl)C12CCC2 |
| Ligand75_54.sdf | CCCCNCc1cn(-c2c(Cl)cc3c(c2F)N/C(=N\C)C32CCC2)nn1 |
| Ligand75_55.sdf | C/N=C1\Nc2c(cc(Cl)c(-n3cc(C4CCCN4)nn3)c2F)C12CCC2 |
| Ligand75_56.sdf | CCCCCCNCC#Cc1cc2c(cc1Cl)C1(CCC1)/C(=N/C)N2 |
| Ligand75_57.sdf | C/N=C1\Nc2cc(N3N=CC4CN(C)CCCC43)c(Cl)cc2C12CCC2 |
| Ligand75_58.sdf | C/N=C1\Nc2cc(-n3cc([C@H]4CCCN4C(=O)OC(C)(C)C)nn3)c(Cl)cc2C12CCC2 |
| Ligand75_59.sdf | C/N=C1\Nc2c(cc(Cl)c(N3N=CC([C@H]4CCCN4)C3C)c2Cl)C12CCC2 |
| Ligand75_6.sdf | FC(F)(F)c1ccc(N2CCN(CCNCc3nc(C4(c5cccc(Cl)c5)CCC4)no3)CC2)nc1 |
| Ligand75_60.sdf | C/N=C1\Nc2cc(-n3cc(C4CCCN4C)nn3)c(Cl)cc2C12CCC2 |
| Ligand75_61.sdf | CN(C)C1=Nc2cc(-n3cc([C@H]4CCCN4)nn3)c(Cl)cc2C12CCC2 |
| Ligand75_62.sdf | C=CNc1c(CC)cc2c(c1Cl)N/C(=N\C)C21CCC1.NCC1CCCN1 |
| Ligand75_63.sdf | C=CNc1c(CC)cc2c(c1Cl)N/C(=N\C)C21CCC1 |
| Ligand75_64.sdf | CCOC(=O)c1cnn(-c2cc3c(cc2Cl)C2(CCC2)/C(=N/C)N3)c1 |
| Ligand75_65.sdf | C/N=C1\Nc2cc(-n3cc(CN4CCC4)cn3)c(Cl)cc2C12CCC2 |
| Ligand75_66.sdf | C/N=C1\Nc2cc(-n3cc(CCl)cn3)c(Cl)cc2C12CCC2 |
| Ligand75_67.sdf | C/N=C1\Nc2cc(B(O)O)c(Cl)cc2C12CCC2 |
| Ligand75_68.sdf | CN(Cc1cn(-c2cc3c(cc2Cl)C2(CCC2)C(N)=N3)nn1)C(=O)OC(C)(C)C |
| Ligand75_69.sdf | C/N=C1\Nc2cc(-c3cc4n(n3)CCN(C(=O)OC(C)(C)C)C4)c(Cl)cc2C12CCC2 |
| Ligand75_7.sdf | ClCc1nc(C2(c3ccc(Cl)c(Cl)c3)CCC2)no1 |
| Ligand75_70.sdf | C/N=C1\Nc2cc(-n3cc(CN(C)C(=O)OC(C)(C)C)nn3)c(Cl)cc2C12CCC2 |
| Ligand75_71.sdf | C/N=C1\Nc2cc(-n3cc(CNC)nn3)c(Cl)cc2C12CCC2 |
| Ligand75_72.sdf | C/N=C1\Nc2cc(-n3cc(CO)cn3)c(Cl)cc2C12CCC2 |
| Ligand75_73.sdf | CNCc1cn(-c2cc3c(cc2Cl)C2(CCC2)C(N)=N3)nn1 |
| Ligand75_74.sdf | C/N=C1\Nc2cc(-n3cc([C@@H]4CCCN4C(=O)OC(C)(C)C)nn3)c(Cl)cc2C12CCC2 |
| Ligand75_75.sdf | Cn1ccc(=O)c(-c2nc(C3(c4cccc(Cl)c4)CCC3)no2)c1 |
| Ligand75_76.sdf | Cn1nnnc1C1(c2cccc(Cl)c2)CCC1 |
| Ligand75_8.sdf | N=C(N)C1(c2cccc(Cl)c2)CCC1 |
| Ligand75_9.sdf | Clc1cccc(C2(C3=NCCN3)CCC2)c1 |
| Ligand_2.sdf | CC(C)(O)c1ccc(-c2cc(Cl)cc(C3(C(=N)N)CCC3)c2)cc1 |
| Ligand_3.sdf | CC(C)(O)c1ccc(-c2cc(Cl)cc(C3(C(=N)N)CCC3)c2)cc1 |
| Ligand_75.sdf | Clc1ccc(C2(c3nnc4n3CCCCCC4)CCC2)cc1Cl |
| Ligand_75.sdf | FC(F)(F)c1cccc(CN2CCN(CCNCc3nc(C4(c5cccc(Cl)c5)CCC4)no3)CC2)c1 |
| Ligand_75.sdf | FC(F)(F)c1cccc(C2CCN(CCNCc3nc(C4(c5cccc(Cl)c5)CCC4)no3)CC2)c1 |
| Ligand_75.sdf | COc1ccccc1C1CCN(CCNCc2nc(C3(c4ccc(Cl)c(Cl)c4)CCC3)no2)CC1 |
| Ligand_75.sdf | FC(F)(F)c1cccc(C2CCN(CCNCc3nc(C4(c5ccc(Cl)c(Cl)c5)CCC4)no3)CC2)c1 |
| Ligand_75.sdf | FC(F)(F)c1ccc(N2CCN(CCNCc3nc(C4(c5cccc(Cl)c5)CCC4)no3)CC2)nc1 |
| Ligand_75.sdf | ClCc1nc(C2(c3ccc(Cl)c(Cl)c3)CCC2)no1 |
| Ligand_75.sdf | N=C(N)C1(c2cccc(Cl)c2)CCC1 |
| Ligand_75.sdf | Clc1cccc(C2(C3=NCCN3)CCC2)c1 |
| Ligand_75.sdf | C/N=C1\Nc2cc(C#CCN3CCCCC3)c(Cl)cc2C12CCC2 |
| Ligand_75.sdf | O=C(Nc1ccc(C2(c3nc4ncc(Br)cc4[nH]3)CCC2)cc1Cl)c1cccc(Br)c1 |
| Ligand_75.sdf | N#Cc1cccc(C(=O)Nc2ccc(C3(c4nc5ncc(C#N)cc5[nH]4)CCC3)cc2Cl)c1 |
| Ligand_75.sdf | CC(C)(O)c1ccc(-c2cc(Cl)cc(C3(C(=N)N)CCC3)c2)cc1 |
| Ligand_75.sdf | CN(Cc1nc(C2(c3cccc(Cl)c3)CCC2)no1)S(C)(=O)=O |
| Ligand_75.sdf | CCC(=O)N1CCCCC1c1nc(C2(c3cccc(Cl)c3)CCC2)no1 |
| Ligand_75.sdf | CCC(=O)N1CCCC(c2nc(C3(c4cccc(Cl)c4)CCC3)no2)C1 |
| Ligand_75.sdf | O=C1CCCN1CCc1nc(C2(c3cccc(Cl)c3)CCC2)no1 |
| Ligand_75.sdf | CC(CS(C)(=O)=O)c1nc(C2(c3cccc(Cl)c3)CCC2)no1 |
| Ligand_75.sdf | CC(C)(c1nc(C2(c3cccc(Cl)c3)CCC2)no1)S(C)(=O)=O |
| Ligand_75.sdf | C/N=C1\Nc2cc(-n3cc([C@H]4CCCN4)nn3)c(Cl)cc2C12CCC2 |
| Ligand_75.sdf | C/N=C1\Nc2cc(-c3cc4n(n3)CCNC4)c(Cl)cc2C12CCC2 |
| Ligand_75.sdf | N=C(N)C1(c2cc(Cl)cc(Br)c2)CCC1 |
| Ligand_75.sdf | NC(=O)c1nc(C2(c3cccc(Cl)c3)CCC2)no1 |
| Ligand_75.sdf | C/N=C1\Nc2c(cc(Cl)c(-n3cc(C4CCCN4)cn3)c2C)C12CCC2 |
| Ligand_75.sdf | C/N=C1\Nc2c(cc(Cl)c(-n3cc(CNC)cn3)c2Cl)C12CCC2 |
| Ligand_75.sdf | C/N=C1\Nc2cc(N(N)/C=C(\N)C3CCCCN3)c(Cl)cc2C12CCC2 |
| Ligand_75.sdf | C/N=C1\Nc2cc(N(N)/C=C(\N)C3CCCN3C)c(Cl)cc2C12CCC2 |
| Ligand_75.sdf | C/N=C1\Nc2c(cc(C)c(C)c2Cl)C12CCC2 |
| Ligand_75.sdf | C/N=C1\Nc2cc(-n3ncc4c3CCN(C)C4)c(Cl)cc2C12CCC2 |
| Ligand_75.sdf | CC1(c2nc(C3(c4cccc(Cl)c4)CCC3)no2)CCNC(=O)C1 |
| Ligand_75.sdf | Cc1cc(Cc2nc(C3(c4cccc(Cl)c4)CCC3)no2)no1 |
| Ligand_75.sdf | C/N=C1\Nc2cc(-n3cc(CN(C)C)cn3)c(Cl)cc2C12CCC2 |
| Ligand_75.sdf | C/N=C1\Nc2c(cc(Cl)c(-n3cc(C4CCCN4)cn3)c2Cl)C12CCC2 |
| Ligand_75.sdf | C/N=C1\Nc2c(cc(Cl)c(-n3cc([C@H]4CCCN4)cn3)c2Cl)C12CCC2 |
| Ligand_75.sdf | C/N=C1\Nc2cc(-n3cc([C@H]4CCCN4)cn3)c(Cl)cc2C12CCC2 |
| Ligand_75.sdf | C/N=C1\Nc2cc(-n3cc([C@@H]4CCCN4)cn3)c(Cl)cc2C12CCC2 |
| Ligand_75.sdf | C/N=C1\Nc2c(cc(Cl)c(-n3cc([C@@H]4CCCN4)cn3)c2Cl)C12CCC2 |
| Ligand_75.sdf | C/N=C1\Nc2c(cc(Cl)c(-n3cccn3)c2Cl)C12CCC2 |
| Ligand_75.sdf | C/N=C1\Nc2c(cc(OC)c(-n3cc(CNC)cn3)c2Cl)C12CCC2 |
| Ligand_75.sdf | C/N=C1\Nc2cc(-n3cc(C4CCCN4)cn3)c(Cl)cc2C12CCC2 |
| Ligand_75.sdf | C/N=C1\Nc2cc(-n3cc(CNC)cn3)c(Cl)cc2C12CCC2 |
| Ligand_75.sdf | C/N=C1\Nc2c(cc(Cl)c(-n3cc(CNC)cn3)c2F)C12CCC2 |
| Ligand_75.sdf | C/N=C1\Nc2c(cc(Cl)c(N3CC([C@@H]4CCCN4)C=N3)c2Cl)C12CCC2 |
| Ligand_75.sdf | C/N=C1\Nc2c(cc(Cl)c(-n3cc(C)cn3)c2Cl)C12CCC2 |
| Ligand_75.sdf | C/N=C1\Nc2cc(-n3cccn3)c(Cl)cc2C12CCC2 |
| Ligand_75.sdf | C/N=C1\Nc2cc(-n3cc(C4CCCN4)nn3)c(Cl)cc2C12CCC2 |
| Ligand_75.sdf | C/N=C1\Nc2cc(-n3cc([C@@H]4CCCN4)nn3)c(Cl)cc2C12CCC2 |
| Ligand_75.sdf | C/N=C1\Nc2cc(-n3cc4c(n3)CCCNC4)c(Cl)cc2C12CCC2 |
| Ligand_75.sdf | C/N=C1\Nc2c(cc(OC)c(-n3cc([C@@H]4CCCN4)nn3)c2Cl)C12CCC2 |
| Ligand_75.sdf | C/N=C1\Nc2cc(-n3cc([C@@H]4CCCCN4)nn3)c(Cl)cc2C12CCC2 |
| Ligand_75.sdf | C/N=C1\Nc2cc(-n3cc(C4CCCCN4)nn3)c(Cl)cc2C12CCC2 |
| Ligand_75.sdf | C/N=C1\Nc2cc(-n3ncc4c3CCNC4)c(Cl)cc2C12CCC2 |
| Ligand_75.sdf | C/N=C1\Nc2c(cc(Cl)c(-n3cc(C4CCCC4)cn3)c2Cl)C12CCC2 |
| Ligand_75.sdf | CCCCNCc1cn(-c2c(Cl)cc3c(c2F)N/C(=N\C)C32CCC2)nn1 |
| Ligand_75.sdf | C/N=C1\Nc2c(cc(Cl)c(-n3cc(C4CCCN4)nn3)c2F)C12CCC2 |
| Ligand_75.sdf | CCCCCCNCC#Cc1cc2c(cc1Cl)C1(CCC1)/C(=N/C)N2 |
| Ligand_75.sdf | C/N=C1\Nc2cc(N3N=CC4CN(C)CCCC43)c(Cl)cc2C12CCC2 |
| Ligand_75.sdf | C/N=C1\Nc2cc(-n3cc([C@H]4CCCN4C(=O)OC(C)(C)C)nn3)c(Cl)cc2C12CCC2 |
| Ligand_75.sdf | C/N=C1\Nc2c(cc(Cl)c(N3N=CC([C@H]4CCCN4)C3C)c2Cl)C12CCC2 |
| Ligand_75.sdf | C/N=C1\Nc2cc(-n3cc(C4CCCN4C)nn3)c(Cl)cc2C12CCC2 |
| Ligand_75.sdf | CN(C)C1=Nc2cc(-n3cc([C@H]4CCCN4)nn3)c(Cl)cc2C12CCC2 |
| Ligand_75.sdf | C=CNc1c(CC)cc2c(c1Cl)N/C(=N\C)C21CCC1.NCC1CCCN1 |
| Ligand_75.sdf | C=CNc1c(CC)cc2c(c1Cl)N/C(=N\C)C21CCC1 |
| Ligand_75.sdf | CCOC(=O)c1cnn(-c2cc3c(cc2Cl)C2(CCC2)/C(=N/C)N3)c1 |
| Ligand_75.sdf | C/N=C1\Nc2cc(-n3cc(CN4CCC4)cn3)c(Cl)cc2C12CCC2 |
| Ligand_75.sdf | C/N=C1\Nc2cc(-n3cc(CCl)cn3)c(Cl)cc2C12CCC2 |
| Ligand_75.sdf | C/N=C1\Nc2cc(B(O)O)c(Cl)cc2C12CCC2 |
| Ligand_75.sdf | CN(Cc1cn(-c2cc3c(cc2Cl)C2(CCC2)C(N)=N3)nn1)C(=O)OC(C)(C)C |
| Ligand_75.sdf | C/N=C1\Nc2cc(-c3cc4n(n3)CCN(C(=O)OC(C)(C)C)C4)c(Cl)cc2C12CCC2 |
| Ligand_75.sdf | C/N=C1\Nc2cc(-n3cc(CN(C)C(=O)OC(C)(C)C)nn3)c(Cl)cc2C12CCC2 |
| Ligand_75.sdf | C/N=C1\Nc2cc(-n3cc(CNC)nn3)c(Cl)cc2C12CCC2 |
| Ligand_75.sdf | C/N=C1\Nc2cc(-n3cc(CO)cn3)c(Cl)cc2C12CCC2 |
| Ligand_75.sdf | CNCc1cn(-c2cc3c(cc2Cl)C2(CCC2)C(N)=N3)nn1 |
| Ligand_75.sdf | C/N=C1\Nc2cc(-n3cc([C@@H]4CCCN4C(=O)OC(C)(C)C)nn3)c(Cl)cc2C12CCC2 |
| Ligand_75.sdf | Cn1ccc(=O)c(-c2nc(C3(c4cccc(Cl)c4)CCC3)no2)c1 |
| Ligand_75.sdf | Cn1nnnc1C1(c2cccc(Cl)c2)CCC1 |
| Ligand_Acebutolol_CID_1978.sdf | CCCC(=O)Nc1ccc(OCC(O)CNC(C)C)c(C(C)=O)c1 |
| Ligand_Amlodipine_CID_2162.sdf | CCOC(=O)C1=C(COCCN)NC(C)=C(C(=O)OC)C1c1ccccc1Cl |
| Ligand_Aspirin_CID_2244.sdf | CC(=O)Oc1ccccc1C(=O)O |
| Ligand_Atenolol_CID_2249.sdf | CC(C)NCC(O)COc1ccc(CC(N)=O)cc1 |
| Ligand_Atorvastin_CID_60823.sdf | CC(C)c1c(C(=O)Nc2ccccc2)c(-c2ccccc2)c(-c2ccc(F)cc2)n1CCC(O)CC(O)CC(=O)O |
| Ligand_Betaxolol_hydrochloride_CID_2369.sdf | CC(C)NCC(O)COc1ccc(CCOCC2CC2)cc1 |
| Ligand_Bisoprolol_CID_2405.sdf | CC(C)NCC(O)COc1ccc(COCCOC(C)C)cc1 |
| Ligand_Carvedilol_CID_2585.sdf | COc1ccccc1OCCNCC(O)COc1cccc2[nH]c3ccccc3c12 |
| Ligand_Cerivastatin_CID_446156.sdf | COCc1c(C(C)C)nc(C(C)C)c(/C=C/C(O)CC(O)CC(=O)O)c1-c1ccc(F)cc1 |
| Ligand_Cholestyramine_CID_70695641.sdf | CCc1ccc(C(C)CCc2ccc([N+](C)(C)C)cc2)cc1 |
| Ligand_Clopidogrel.sdf | COC(=O)C(c1ccccc1Cl)N1CCc2sccc2C1 |
| Ligand_Colestipol_CID_8197.sdf | NCCNCCNCCNCCN |
| Ligand_Enalapril_Maleate_CID_5388962.sdf | CCOC(=O)C(CCc1ccccc1)NC(C)C(=O)N1CCCC1C(=O)O |
| Ligand_Esmolol_CID_59768.sdf | COC(=O)CCc1ccc(OCC(O)CNC(C)C)cc1 |
| Ligand_Ezetimibe_CID_150311.sdf | O=C1C(CCC(O)c2ccc(F)cc2)C(c2ccc(O)cc2)N1c1ccc(F)cc1 |
| Ligand_Fluvastatin_CID_446155.sdf | CC(C)n1c(/C=C/C(O)CC(O)CC(=O)O)c(-c2ccc(F)cc2)c2ccccc21 |
| Ligand_Isoxsuprine_CID_3783.sdf | CC(COc1ccccc1)NC(C)C(O)c1ccc(O)cc1 |
| Ligand_Lisinopril_CID_5362119.sdf | NCCCCC(NC(CCc1ccccc1)C(=O)O)C(=O)N1CCCC1C(=O)O |
| Ligand_Lovastatin_CID_53232.sdf | CCC(C)C(=O)OC1CC(C)C=C2C=CC(C)C(CCC3CC(O)CC(=O)O3)C21 |
| Ligand_Metoprolol_CID_4171.sdf | COCCc1ccc(OCC(O)CNC(C)C)cc1 |
| Ligand_Nadolol_CID_39147.sdf | CC(C)(C)NCC(O)COc1cccc2c1CC(O)C(O)C2 |
| Ligand_Nebivolol_CID_71301.sdf | OC(CNCC(O)C1CCc2cc(F)ccc2O1)C1CCc2cc(F)ccc2O1 |
| Ligand_Perindopril_CID_107807.sdf | CCCC(NC(C)C(=O)N1C(C(=O)O)CC2CCCCC21)C(=O)OCC |
| Ligand_Pitavastatin_CID_5282452.sdf | O=C(O)CC(O)CC(O)/C=C/c1c(C2CC2)nc2ccccc2c1-c1ccc(F)cc1 |
| Ligand_Prasugrel.sdf | CC(=O)Oc1cc2c(s1)CCN(C(C(=O)C1CC1)c1ccccc1F)C2 |
| Ligand_Probucol_CID_4912.sdf | CC(C)(Sc1cc(C(C)(C)C)c(O)c(C(C)(C)C)c1)Sc1cc(C(C)(C)C)c(O)c(C(C)(C)C)c1 |
| Ligand_Propranolol_CID_4946.sdf | CC(C)NCC(O)COc1cccc2ccccc12 |
| Ligand_Rivaroxaban_CID_9875401.sdf | O=C(NCC1CN(c2ccc(N3CCOCC3=O)cc2)C(=O)O1)c1ccc(Cl)s1 |
| Ligand_Rosuvastatin_CID_446157.sdf | CC(C)c1nc(N(C)S(C)(=O)=O)nc(-c2ccc(F)cc2)c1/C=C/C(O)CC(O)CC(=O)O |
| Ligand_Simvastatin_CID_54454.sdf | CCC(C)(C)C(=O)OC1CC(C)C=C2C=CC(C)C(CCC3CC(O)CC(=O)O3)C21 |
| Ligand_Ticagrelor.sdf | CCCSc1nc(NC2CC2c2ccc(F)c(F)c2)c2nnn(C3CC(OCCO)C(O)C3O)c2n1 |
| Ligand_2.sdf | CC(C)(O)c1ccc(-c2cc(Cl)cc(C3(C(=N)N)CCC3)c2)cc1 |
| Ligand_3.sdf | CC(C)(O)c1ccc(-c2cc(Cl)cc(C3(C(=N)N)CCC3)c2)cc1 |
| Ligand_75.sdf | Clc1ccc(C2(c3nnc4n3CCCCCC4)CCC2)cc1Cl |
| Ligand_75.sdf | FC(F)(F)c1cccc(CN2CCN(CCNCc3nc(C4(c5cccc(Cl)c5)CCC4)no3)CC2)c1 |
| Ligand_75.sdf | FC(F)(F)c1cccc(C2CCN(CCNCc3nc(C4(c5cccc(Cl)c5)CCC4)no3)CC2)c1 |
| Ligand_75.sdf | COc1ccccc1C1CCN(CCNCc2nc(C3(c4ccc(Cl)c(Cl)c4)CCC3)no2)CC1 |
| Ligand_75.sdf | FC(F)(F)c1cccc(C2CCN(CCNCc3nc(C4(c5ccc(Cl)c(Cl)c5)CCC4)no3)CC2)c1 |
| Ligand_75.sdf | FC(F)(F)c1ccc(N2CCN(CCNCc3nc(C4(c5cccc(Cl)c5)CCC4)no3)CC2)nc1 |
| Ligand_75.sdf | ClCc1nc(C2(c3ccc(Cl)c(Cl)c3)CCC2)no1 |
| Ligand_75.sdf | N=C(N)C1(c2cccc(Cl)c2)CCC1 |
| Ligand_75.sdf | Clc1cccc(C2(C3=NCCN3)CCC2)c1 |
| Ligand_75.sdf | C/N=C1\Nc2cc(C#CCN3CCCCC3)c(Cl)cc2C12CCC2 |
| Ligand_75.sdf | O=C(Nc1ccc(C2(c3nc4ncc(Br)cc4[nH]3)CCC2)cc1Cl)c1cccc(Br)c1 |
| Ligand_75.sdf | N#Cc1cccc(C(=O)Nc2ccc(C3(c4nc5ncc(C#N)cc5[nH]4)CCC3)cc2Cl)c1 |
| Ligand_75.sdf | CC(C)(O)c1ccc(-c2cc(Cl)cc(C3(C(=N)N)CCC3)c2)cc1 |
| Ligand_75.sdf | CN(Cc1nc(C2(c3cccc(Cl)c3)CCC2)no1)S(C)(=O)=O |
| Ligand_75.sdf | CCC(=O)N1CCCCC1c1nc(C2(c3cccc(Cl)c3)CCC2)no1 |
| Ligand_75.sdf | CCC(=O)N1CCCC(c2nc(C3(c4cccc(Cl)c4)CCC3)no2)C1 |
| Ligand_75.sdf | O=C1CCCN1CCc1nc(C2(c3cccc(Cl)c3)CCC2)no1 |
| Ligand_75.sdf | CC(CS(C)(=O)=O)c1nc(C2(c3cccc(Cl)c3)CCC2)no1 |
| Ligand_75.sdf | CC(C)(c1nc(C2(c3cccc(Cl)c3)CCC2)no1)S(C)(=O)=O |
| Ligand_75.sdf | C/N=C1\Nc2cc(-n3cc([C@H]4CCCN4)nn3)c(Cl)cc2C12CCC2 |
| Ligand_75.sdf | C/N=C1\Nc2cc(-c3cc4n(n3)CCNC4)c(Cl)cc2C12CCC2 |
| Ligand_75.sdf | N=C(N)C1(c2cc(Cl)cc(Br)c2)CCC1 |
| Ligand_75.sdf | NC(=O)c1nc(C2(c3cccc(Cl)c3)CCC2)no1 |
| Ligand_75.sdf | C/N=C1\Nc2c(cc(Cl)c(-n3cc(C4CCCN4)cn3)c2C)C12CCC2 |
| Ligand_75.sdf | C/N=C1\Nc2c(cc(Cl)c(-n3cc(CNC)cn3)c2Cl)C12CCC2 |
| Ligand_75.sdf | C/N=C1\Nc2cc(N(N)/C=C(\N)C3CCCCN3)c(Cl)cc2C12CCC2 |
| Ligand_75.sdf | C/N=C1\Nc2cc(N(N)/C=C(\N)C3CCCN3C)c(Cl)cc2C12CCC2 |
| Ligand_75.sdf | C/N=C1\Nc2c(cc(C)c(C)c2Cl)C12CCC2 |
| Ligand_75.sdf | C/N=C1\Nc2cc(-n3ncc4c3CCN(C)C4)c(Cl)cc2C12CCC2 |
| Ligand_75.sdf | CC1(c2nc(C3(c4cccc(Cl)c4)CCC3)no2)CCNC(=O)C1 |
| Ligand_75.sdf | Cc1cc(Cc2nc(C3(c4cccc(Cl)c4)CCC3)no2)no1 |
| Ligand_75.sdf | C/N=C1\Nc2cc(-n3cc(CN(C)C)cn3)c(Cl)cc2C12CCC2 |
| Ligand_75.sdf | C/N=C1\Nc2c(cc(Cl)c(-n3cc(C4CCCN4)cn3)c2Cl)C12CCC2 |
| Ligand_75.sdf | C/N=C1\Nc2c(cc(Cl)c(-n3cc([C@H]4CCCN4)cn3)c2Cl)C12CCC2 |
| Ligand_75.sdf | C/N=C1\Nc2cc(-n3cc([C@@H]4CCCN4)cn3)c(Cl)cc2C12CCC2 |
| Ligand_75.sdf | C/N=C1\Nc2cc(-n3cc([C@@H]4CCCN4)cn3)c(Cl)cc2C12CCC2 |
| Ligand_75.sdf | C/N=C1\Nc2c(cc(Cl)c(-n3cc([C@H]4CCCN4)cn3)c2Cl)C12CCC2 |
| Ligand_75.sdf | C/N=C1\Nc2c(cc(Cl)c(-n3cccn3)c2Cl)C12CCC2 |
| Ligand_75.sdf | C/N=C1\Nc2c(cc(OC)c(-n3cc(CNC)cn3)c2Cl)C12CCC2 |
| Ligand_75.sdf | C/N=C1\Nc2cc(-n3cc(C4CCCN4)cn3)c(Cl)cc2C12CCC2 |
| Ligand_75.sdf | C/N=C1\Nc2cc(-n3cc(CNC)cn3)c(Cl)cc2C12CCC2 |
| Ligand_75.sdf | C/N=C1\Nc2c(cc(Cl)c(-n3cc(CNC)cn3)c2F)C12CCC2 |
| Ligand_75.sdf | C/N=C1\Nc2c(cc(Cl)c(N3C[C@H]([C@H]4CCCN4)C=N3)c2Cl)C12CCC2 |
| Ligand_75.sdf | C/N=C1\Nc2c(cc(Cl)c(-n3cc(C)cn3)c2Cl)C12CCC2 |
| Ligand_75.sdf | C/N=C1\Nc2cc(-n3cccn3)c(Cl)cc2C12CCC2 |
| Ligand_75.sdf | C/N=C1\Nc2cc(-n3cc(C4CCCN4)nn3)c(Cl)cc2C12CCC2 |
| Ligand_75.sdf | C/N=C1\Nc2cc(-n3cc([C@H]4CCCN4)nn3)c(Cl)cc2C12CCC2 |
| Ligand_75.sdf | C/N=C1\Nc2cc(-n3cc4c(n3)CCCNC4)c(Cl)cc2C12CCC2 |
| Ligand_75.sdf | C/N=C1\Nc2c(cc(OC)c(-n3cc([C@H]4CCCN4)nn3)c2Cl)C12CCC2 |
| Ligand_75.sdf | C/N=C1\Nc2cc(-n3cc([C@H]4CCCCN4)nn3)c(Cl)cc2C12CCC2 |
| Ligand_75.sdf | C/N=C1\Nc2cc(-n3cc(C4CCCCN4)nn3)c(Cl)cc2C12CCC2 |
| Ligand_75.sdf | C/N=C1\Nc2cc(-n3ncc4c3CCNC4)c(Cl)cc2C12CCC2 |
| Ligand_75.sdf | C/N=C1/Nc2c(cc(Cl)c(-n3cc(C4CCCC4)cn3)c2Cl)C12CCC2 |
| Ligand_75.sdf | CCCCNCc1cn(-c2c(Cl)cc3c(c2F)N/C(=N\C)C32CCC2)nn1 |
| Ligand_75.sdf | C/N=C1\Nc2c(cc(Cl)c(-n3cc(C4CCCN4)nn3)c2F)C12CCC2 |
| Ligand_75.sdf | CCCCCCNCC#Cc1cc2c(cc1Cl)C1(CCC1)/C(=N/C)N2 |
| Ligand_75.sdf | C/N=C1\Nc2cc(N3N=CC4CN(C)CCCC43)c(Cl)cc2C12CCC2 |
| Ligand_75.sdf | C/N=C1\Nc2cc(-n3cc([C@H]4CCCN4C(=O)OC(C)(C)C)nn3)c(Cl)cc2C12CCC2 |
| Ligand_75.sdf | C/N=C1\Nc2c(cc(Cl)c(N3N=C[C@@H]([C@H]4CCCN4)[C@@H]3C)c2Cl)C12CCC2 |
| Ligand_75.sdf | C/N=C1\Nc2cc(-n3cc(C4CCCN4C)nn3)c(Cl)cc2C12CCC2 |
| Ligand_75.sdf | CN(C)C1=Nc2cc(-n3cc([C@H]4CCCN4)nn3)c(Cl)cc2C12CCC2 |
| Ligand_75.sdf | C=CNc1c(CC)cc2c(c1Cl)N/C(=N\C)C21CCC1.NCC1CCCN1 |
| Ligand_75.sdf | C=CNc1c(CC)cc2c(c1Cl)N/C(=N\C)C21CCC1 |
| Ligand_75.sdf | CCOC(=O)c1cnn(-c2cc3c(cc2Cl)C2(CCC2)/C(=N/C)N3)c1 |
| Ligand_75.sdf | C/N=C1\Nc2cc(-n3cc(CN4CCC4)cn3)c(Cl)cc2C12CCC2 |
| Ligand_75.sdf | C/N=C1\Nc2cc(-n3cc(CCl)cn3)c(Cl)cc2C12CCC2 |
| Ligand_Acebutolol_CID_1978.sdf | CCCC(=O)Nc1ccc(OC[C@H](O)CNC(C)C)c(C(C)=O)c1 |
| Ligand_Amlodipine_CID_2162.sdf | CCOC(=O)C1=C(COCCN)NC(C)=C(C(=O)OC)[C@H]1c1ccccc1Cl |
| Ligand_Aspirin_CID_2244.sdf | CC(=O)Oc1ccccc1C(=O)O |
| Ligand_Atenolol_CID_2249.sdf | CC(C)NC[C@@H](O)COc1ccc(CC(N)=O)cc1 |
| Ligand_Atorvastin_CID_60823.sdf | CC(C)c1c(C(=O)Nc2ccccc2)c(-c2ccccc2)c(-c2ccc(F)cc2)n1CC[C@@H](O)C[C@@H](O)CC(=O)O |
| Ligand_Betaxolol_hydrochloride_CID_2369.sdf | CC(C)NC[C@@H](O)COc1ccc(CCOCC2CC2)cc1 |
| Ligand_Bisoprolol_CID_2405.sdf | CC(C)NC[C@@H](O)COc1ccc(COCCOC(C)C)cc1 |
| Ligand_Carvedilol_CID_2585.sdf | COc1ccccc1OCCNC[C@@H](O)COc1cccc2[nH]c3ccccc3c12 |
| Ligand_Cerivastatin_CID_446156.sdf | COCc1c(C(C)C)nc(C(C)C)c(/C=C/[C@@H](O)C[C@@H](O)CC(=O)O)c1-c1ccc(F)cc1 |
| Ligand_Cholestyramine_CID_70695641.sdf | CCc1ccc([C@H](C)CCc2ccc([N+](C)(C)C)cc2)cc1 |
| Ligand_Clopidogrel.sdf | COC(=O)[C@H](c1ccccc1Cl)N1CCc2sccc2C1 |
| Ligand_Colestipol_CID_8197.sdf | NCCNCCNCCNCCN |
| Ligand_Enalapril_Maleate_CID_5388962.sdf | CCOC(=O)[C@H](CCc1ccccc1)N[C@@H](C)C(=O)N1CCC[C@H]1C(=O)O |
| Ligand_Esmolol_CID_59768.sdf | COC(=O)CCc1ccc(OC[C@H](O)CNC(C)C)cc1 |
| Ligand_Ezetimibe_CID_150311.sdf | O=C1[C@H](CC[C@H](O)c2ccc(F)cc2)[C@@H](c2ccc(O)cc2)N1c1ccc(F)cc1 |
| Ligand_Fluvastatin_CID_446155.sdf | CC(C)n1c(/C=C/[C@@H](O)C[C@@H](O)CC(=O)O)c(-c2ccc(F)cc2)c2ccccc21 |
| Ligand_Isoxsuprine_CID_3783.sdf | C[C@H](COc1ccccc1)N[C@H](C)[C@H](O)c1ccc(O)cc1 |
| Ligand_Lisinopril_CID_5362119.sdf | NCCCC[C@H](N[C@@H](CCc1ccccc1)C(=O)O)C(=O)N1CCC[C@H]1C(=O)O |
| Ligand_Lovastatin_CID_53232.sdf | CC[C@H](C)C(=O)O[C@H]1C[C@@H](C)C=C2C=C[C@H](C)[C@H](CC[C@@H]3C[C@@H](O)CC(=O)O3)[C@H]21 |
| Ligand_Metoprolol_CID_4171.sdf | COCCc1ccc(OC[C@H](O)CNC(C)C)cc1 |
| Ligand_Nadolol_CID_39147.sdf | CC(C)(C)NC[C@H](O)COc1cccc2c1C[C@H](O)[C@H](O)C2 |
| Ligand_Nebivolol_CID_71301.sdf | O[C@H](CNC[C@@H](O)[C@H]1CCc2cc(F)ccc2O1)[C@H]1CCc2cc(F)ccc2O1 |
| Ligand_Perindopril_CID_107807.sdf | CCC[C@H](N[C@@H](C)C(=O)N1[C@H](C(=O)O)C[C@@H]2CCCC[C@@H]21)C(=O)OCC |
| Ligand_Pitavastatin_CID_5282452.sdf | O=C(O)C[C@H](O)C[C@H](O)/C=C/c1c(C2CC2)nc2ccccc2c1-c1ccc(F)cc1 |
| Ligand_Prasugrel.sdf | CC(=O)Oc1cc2c(s1)CCN([C@@H](C(=O)C1CC1)c1ccccc1F)C2 |
| Ligand_Probucol_CID_4912.sdf | CC(C)(Sc1cc(C(C)(C)C)c(O)c(C(C)(C)C)c1)Sc1cc(C(C)(C)C)c(O)c(C(C)(C)C)c1 |
| Ligand_Propranolol_CID_4946.sdf | CC(C)NC[C@H](O)COc1cccc2ccccc12 |
| Ligand_Rivaroxaban_CID_9875401.sdf | O=C(NC[C@H]1CN(c2ccc(N3CCOCC3=O)cc2)C(=O)O1)c1ccc(Cl)s1 |
| Ligand_Rosuvastatin_CID_446157.sdf | CC(C)c1nc(N(C)S(C)(=O)=O)nc(-c2ccc(F)cc2)c1/C=C/[C@@H](O)C[C@@H](O)CC(=O)O |
| Ligand_Simvastatin_CID_54454.sdf | CCC(C)(C)C(=O)O[C@H]1C[C@@H](C)C=C2C=C[C@H](C)[C@H](CC[C@@H]3C[C@@H](O)CC(=O)O3)[C@H]21 |
| Ligand_Ticagrelor.sdf | CCCSc1nc(N[C@@H]2C[C@H]2c2ccc(F)c(F)c2)c2nnn([C@@H]3C[C@H](OCCO)[C@@H](O)[C@H]3O)c2n1 |
